# Constitutively active RAS in *S. pombe* causes persistent Cdc42 signalling but only transient MAPK activation

**DOI:** 10.1101/380220

**Authors:** Emma J. Kelsall, Ábel Vértesy, Kees Straatman, Mishal Tariq, Raquel Gadea, Chandni Parmar, Gabriele Schreiber, Shubhchintan Randhawa, Cyril Dominguez, Edda Klipp, Kayoko Tanaka

## Abstract

The small GTPase RAS is a signalling hub for many pathways and oncogenic human RAS mutations are assumed to over-activate all of its downstream pathways. We tested this assumption in fission yeast, where, RAS-mediated pheromone signalling (PS) activates the MAPK^Spk1^ and Cdc42 pathways. Unexpectedly, we found that constitutively active Ras1.G17V induced immediate but only transient MAPK^Spk1^ activation, whilst Cdc42 activation persisted. Immediate but transient MAPK^Spk1^ activation was also seen in the deletion mutant of Cdc42-GEF^Scd1^, a Cdc42 activator. We built a mathematical model using PS negative-feedback circuits and competition between the two Ras1 effectors, MAPKKK^Byr2^ and Cdc42-GEF^Scd1^. The model robustly predicted the MAPK^Spk1^ activation dynamics of an additional 21 PS mutants. Supporting the model, we showed that a recombinant Cdc42-GEF^Scd1^ fragment competes with MAPKKK^Byr2^ for Ras1 binding. Our study has established a concept that the constitutively active RAS propagates differently to downstream pathways where the system prevents MAPK overactivation.

**Highlights:**

1. Constitutively active Ras1.GV prolongs Cdc42 activation in *S. pombe* pheromone signalling
2. Ras1.GV results in an immediate but only transient MAPK^Spk1^ activation
3. The RAS effector pathways MAPK^Spk1^ and Cdc42 compete with each other for active Ras1
4. Predictive modelling explains MAPK^Spk1^ activation dynamics in 24 signaling-mutants

**eTOC Blurb:** *S. pombe* Ras1 activates the MAPK^Spk1^ and Cdc42 pathways. Kelsall et al. report that the constitutively active Ras1.G17V mutation, which causes morphological anomalies, induces prolonged Cdc42 activation but only a transient MAPK^Spk1^ activation followed by attenuation. Mathematical modelling and biochemical data suggest a competition between the MAPK^Spk1^ and Cdc42 pathways for active Ras1.

## Introduction

Proto-oncogene Ras GTPase family members are widely conserved and play pivotal roles in cell growth, differentiation and apoptosis (Cox and Der, 2010). The physiological impact of Ras mutations is highlighted in the resultant tumorigenesis and developmental disorders (Prior et al., 2012; Schubbert et al., 2007). More than 99% of identified oncogenic RAS mutations occur at codons 12, 13 and 61 of human Ras isoforms (Prior et al., 2012) and impair efficient GTP hydrolysis (Trahey and McCormick, 1987). This results in accumulation of GTP-bound Ras, which is generally considered to cause constitutive activation of the downstream effector pathways, such as ERK and PI3K signalling pathways (Lin et al., 1998; Zhu et al., 1998). Interestingly, however, mouse embryonic fibroblasts (MEFs) derived from the *K-ras^G12D^* mouse model show neither an increased basal level of active ERK and Akt nor constitutive activation of ERK and Akt upon growth factor stimulation even though the *K-ras^G12D^* MEFs showed enhanced proliferation and partial transformation (Tuveson et al., 2004). This observation indicates that not all effector pathways become constitutively activated by oncogenic Ras, presumably because of an efficient negative feedback loop. Meanwhile, it is reasonable to expect that oncogenic RAS over-activates some of its effector pathways to initiate tumorigenesis. Small GTPases, including Cdc42 and Rac, may be such effector pathways, as they are required in oncogenic-RAS-driven tumorigenesis (Malliri et al., 2002; Qiu et al., 1997; Stengel and Zheng, 2012).

We wished to understand how multiple Ras effector pathways respond to the Ras-triggered signals in a physiological setting. We employed the model organism fission yeast. In this organism, a unique Ras homologue, Ras1, plays a key role in pheromone signalling to cause mating of haploid cells and sporulation in diploid cells (Yamamoto, 1996)(Fig. 1A, B). Upon nutritional starvation, cells of opposite mating types (*h^+^* and *h^−^*) exchange mating pheromones. Gpa1, the α-subunit of the pheromone receptor-coupled G-protein, relays the pheromone signal into the cell (Obara et al., 1991). Activated Gpa1 together with Ras1 then activate MAPK cascade consisting of Byr2 (MAPKKK), Byr1 (MAPKK) and Spk1 (MAPK) (Fukui et al., 1986; Masuda et al., 1995; Nadin-Davis and Nasim, 1988; Nadin-Davis et al., 1986a; Nadin-Davis et al., 1986b; Wang et al., 1991; Xu et al., 1994). An intriguing observation is that the *ras1.G17V* mutant, an equivalent of mammalian *ras.G12V* mutant prevalent in cancer, produces an excessively elongated shmoo, or a conjugation tube, upon exposure to the mating pheromone (Nadin-Davis et al., 1986a). This “elongated” *ras1.G17V* phenotype has been interpretted that Ras1 may be responsible for amplifying the pheromone signal (Yamamoto, 1996).

**Figure 1.**
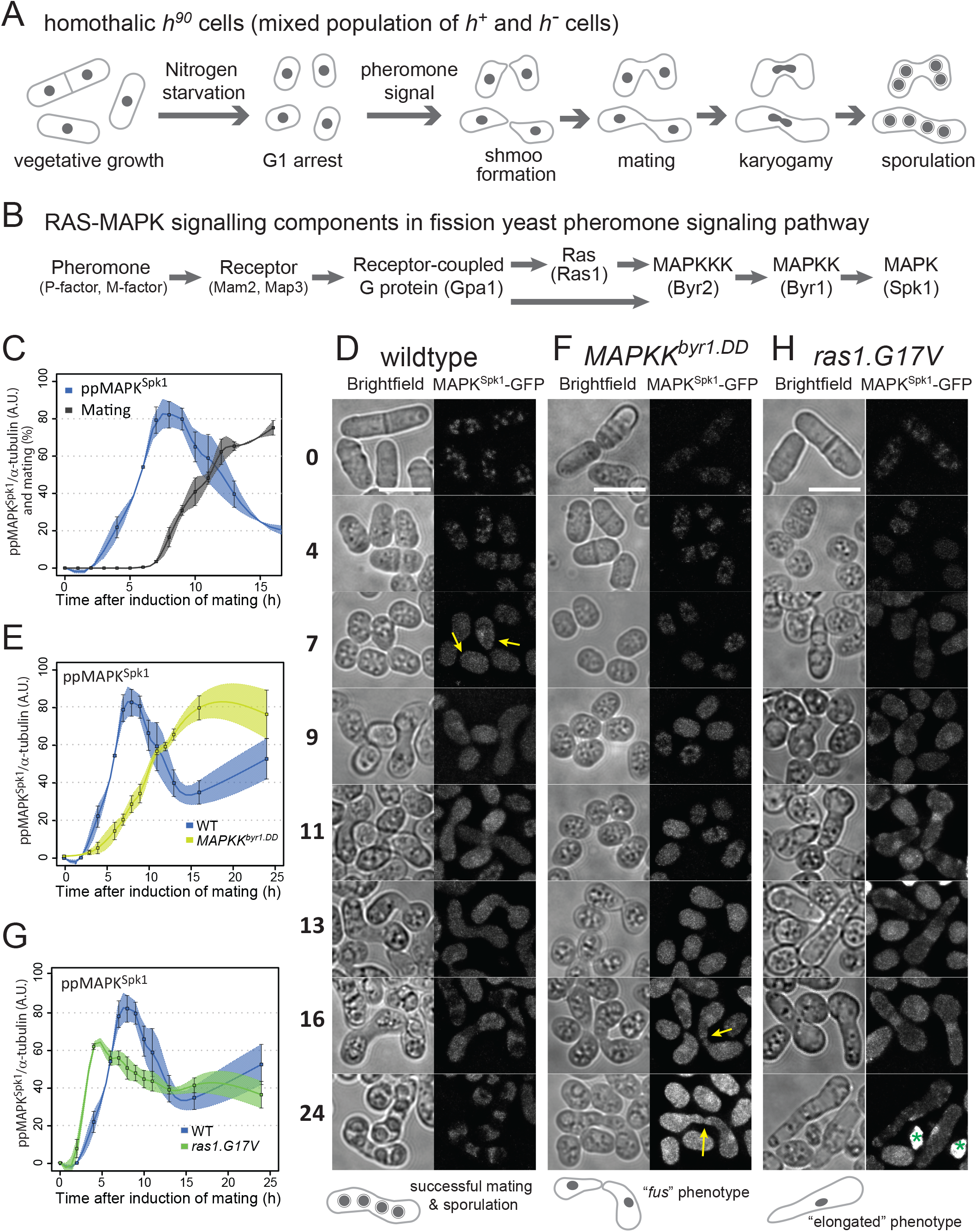
Distinct modes of MAPK^Spk1^ temporal phosphorylation profile and morphological changes during sexual differentiation in wildtype, *MAPKK^byr1.DD^* and *ras1.G17V* mutants. (A) A pictorial representation of wildtype fission yeast sexual differentiation. (B) A list of key signalling components of the fission yeast pheromone signalling pathway. The diagram reflects the prediction that Gpa1 and Ras1 separately contribute to activation of MAPKKK^Byr2^ activation although the precise mechanism is unknown (Xu et al., 1994). At the same time, Ras1 activation is expected to be at least partly under influence of active Gpa1 because the *ste6* gene, encoding a Ras1 activator, is strongly induced upon successful pheromone signaling (Hughes et al., 1994). (C)-(H) Cells were induced for sexual differentiation by the plate mating assay system as described in the materials and methods. (C), (E) and (G) Quantified **pp**MAPK^Spk1^ signal from western blots of wildtype (KT3082) (C), *MAPKK^byr1.DD^* (KT3435) (E) and *ras1.G17V*(KT3084) (G) cells. Three biological replicates were used for quantitation (error bars are ±SEM). α-tubulin was used as a loading control and quantitation was carried out using the Image Studio ver2.1 software (Licor Odyssey CLx Scanner). For the wildtype samples in (C), the % of cells mating is also indicated (n=400, three biological replicates). The wildtype **pp**MAPK^Spk1^ result (C) is also presented in (E) and (G) as a reference. (D), (F) and (H) Cellular morphology (brightfield) and localization of MAPK^Spk1^-GFP over a 24 hour time-course in wildtype (D), *MAPKK^byr1.DD^* (F) and *ras1.G17V* (H) cells. Time after induction of mating in hours is indicated on the left. At each time point, a bright-field image and a GFP signal image were taken and processed as described in materials and methods. Green asterisks in the time 24 h in the *ras1.G17V* cell image (H) indicate auto-fluorescence signal from inviable cell debris, which were produced through cytokinesis failure or cell lysis. Yellow arrows in panels (D) and (F) indicate transient accumulation of MAPK^Spk1^-GFP at the shmoo tips. Scale bars represent 10μm.

Ras1 also regulates cell morphology during vegetative growth; whilst deletion of either *gpa1, MAPKKK^byr2^, MAPKK^byr1^* or *MAPK^spk1^* does not result in any obvious phenotypes during vegetative cell growth (Obara et al., 1991; Sipiczki, 1988; Toda et al., 1991), *ras1Δ* cells lose the typical rod-shape morphology of fission yeast to become rounded (Fukui et al., 1989; Nadin-Davis et al., 1986a). Studies based on recombinant protein assays and yeast-2-hybrid analysis demonstrated that Ras1 interacts with both MAPKKK^Byr2^ and Scd1, a GDP-GTP exchange factor (GEF) for Cdc42, which regulates the actin cytoskeleton and cell morphology (Chang et al., 1994; Gronwald et al., 2001; Tu et al., 1997). These observations suggest that Ras1 simultaneously regulates both the pheromone MAPK^Spk1^ and the Cdc42 pathways at the cell membrane (Weston et al., 2013). Indeed, a dynamic “Cdc42 zone” at the cell cortex prior to mating has been observed (Merlini et al., 2013) and Ras1 and MAPK^Spk1^ cascade components are found there and are involved in the process (Dudin et al., 2016; Merlini et al., 2013; Merlini et al., 2016; Merlini et al., 2018; Weston et al., 2013). However, it is not yet understood how Ras1 interplay between the two pathways. By establishing conditions to induce highly synchronous mating of fission yeast cells, for the first time we were able to follow MAPK^Spk1^ activation dynamics during the physiological mating process in wildtype and in mutants showing various mating phenotypes. This comprehensive set of quantitative measurements allowed us to build a mathematical model of the Ras-mediated pheromone signalling. Our model serves as a prototype of a branched Ras-mediated signalling pathway, demonstrating a competition of downstream pathways for a common upstream activator, Ras1. The model also highlights the physiological importance of the bipartite activation of MAPKKK^Byr2^: a Ras1-dependent and a Ras1-independent mechanism, the latter of which employs the adaptor protein Ste4 (Barr et al., 1996; Okazaki et al., 1991). The adaptor^Ste4^ can be targeted to downregulate MAPK^Spk1^ even in the presence of *ras1.G17V* mutation. Finally, our study reveals the crucial role played by Cdc42 in the *ras1.G17V* mutant causing the *ras1.G17V* phenotype.

## Results

### (1) A highly synchronous mating assay allows the precise measurement of MAPK^Spk1^ activity

To directly measure fission yeast pheromone signalling, we quantitated MAPK^Spk1^ phosphorylation throughout the mating process (Supplementary Fig. S1, Fig. 1A and B). We established a protocol to induce highly synchronous mating and employed cells where the endogenous MAPK^Spk1^ is tagged with GFP-2xFLAG. Under these conditions, homothallic *h^90^* cells started to mate 7-hours after induction of mating (Fig. 1C, grey line). Phosphorylated (active) MAPK^Spk1^ (**pp**MAPK^Spk1^) levels were quantitated as described in Materials and Methods. The **pp**MAPK^Spk1^ signal was first detected three hours after induction of mating and reached its peak at about seven hours, when cell fusion was also initially observed (Fig. 1C, blue line. Original membrane images presented in Supplementary Fig. S3). The **pp**MAPK^Spk1^ then gradually decreased to a non-zero level as meiosis continued towards sporulation. The observation established that the MAPK^Spk1^-GFP activation occurs as the mating process progresses and declines before sporulation, around 15 hours post induction. It was also noted that the total MAPK^Spk1^-GFP was essentially not expressed during the vegetative cycle, but it was promptly induced by nitrogen starvation (Supplementary Fig. S2A and S2B). The *mapk^spk1^* gene is a known target of the transcription factor Ste11 (Mata and Bahler, 2006), which itself is activated (phosphorylated) by MAPK^Spk1^ (Kjaerulff et al., 2005). This positive feedback loop likely facilitates a swift increase of MAPK^Spk1^ expression upon nitrogen starvation. We found that MAPK^Spk1^-GFP localised to both the cytosol and the nucleus, with some nuclear accumulation, before it gradually disappeared as the mating process came to the end (Fig. 1D). Interestingly, transient foci of GFP signals were also found at the cell cortex, as has been reported for MAPKK^Byr1^, the activator of MAPK^Spk1^ (Dudin et al., 2016) (Fig. 1D, yellow arrows).

### (2) Constitutively active MAPKK^Byr1.DD^ mutant causes constitutive activation of MAPK^Spk1^

Activation of MAPKK family kinases is mediated by dual phosphorylation of conserved Ser/Thr residues (Zheng and Guan, 1994). These correspond to serine 214 and threonine 218 of the MAPKK^Byr1^. A MAPKK^Byr1^ mutant termed MAPKK^Byr1.DD^, which carries aspartic acid substitution at these sites, was expected to act as a constitutively active MAPKK (Ozoe et al., 2002). We introduced the *MAPKK^byr1.DD^* mutation at its chromosome locus and measured the activation profile of its target, MAPK^Spk1^ during the mating process. The increase of the **pp**MAPK^Spk1^ signal in the *MAPKK^byr1.DD^* mutant strain was delayed compared to the wildtype strain and the **pp**MAPK^Spk1^ accumulated at a slower rate (Fig. 1E, light green line, original membrane images in Fig. S3). However, the level of **pp**MAPK^Spk1^ remained high after reaching its highest intensity at around 16 hours after induction, resulting in a constitutive phosphorylation of MAPK^Spk1^ (Fig. 1E and Supplementary Fig. S2C and S2D). This result highlights that the suggested dephosphorylation of **pp**MAPK^Spk1^ by phosphatases Pmp1 and Pyp1 (Didmon et al., 2002) is not efficient in downregulating the pheromone signalling in the presence of MAPKK^Byr1.DD^. The MAPKK^Byr1.DD^ strain is also less competent in finding the partner mating cell than cells with wildtype MAPKK^Byr1^. We also confirmed the intriguing *“fus”* (fusion deficient) phenotype of cells expressing MAPKK^Byr1.DD^ from its native promoter (Fig.1F) (Dudin et al., 2016; Ozoe et al., 2002). These cells find their partners and pair up successfully but fail to fuse with each other, resulting in *fus* phenotype. Consistent with the observed slower increase in **pp**MAPK^Spk1^ (Fig.1E) the nuclear localisation of MAPK^Spk1^ was also delayed compared to Wildtype cells (Fig 1F). A strong MAPK^Spk1^-GFP signal was then observed in the nuclei of the paired *fus* cells and the nuclear MAPK^Spk1^-GFP signal persists even 24 hours after the induction of mating (Fig. 1F). Interestingly, the projection tips of the paring cells often show increased MAPK^Spk1^-GFP signal (Fig. 1F, yellow arrows). These results conclude that MAPK^Spk1^ activation is highly influenced by the MAPKK^Byr1^ status and phosphatases directly regulating MAPK^Spk1^ cannot counteract MAPKK^Byr1.DD^.

### (3) The *ras1.G17V* mutation causes immediate but transient MAPK^Spk1^ activation

The fission yeast equivalent of human oncogenic *ras.G12V* is *ras1.G17V*, which induces an excessively elongated shmoo, and the cells fail to recognize a partner and become sterile (Fukui et al., 1986; Nadin-Davis et al., 1986a). The “elongated shmoo” phenotype was interpreted as an excess activation of the downstream pathway(s) of Ras1, leading to a prediction that the *ras1.G17V* causes over-activation of MAPK^Spk1^ (Weston et al., 2013; Yamamoto, 1996). However, no direct evidence has been provided.

Quantitation of the **pp**MAPK^Spk1^-GFP in the *ras1.G17V* mutant showed an immediate increase of the **pp**MAPK^Spk1^-GFP upon induction of mating (Fig. 1G, Green line, original membrane images in Fig. S3). However, the signal intensity declined gradually and by 16 hours after induction the level was comparable to wildtype cells, indicating that down-regulation of **pp**MAPK^Spk1-GFP^ is effective, unlike in *MAPKK^byr1.DD^* mutant. Correspondingly, the cellular MAPK^Spk1^-GFP signal also declined 24 hours after induction of mating (Fig. 1H). Collectively, these observations indicate that the down-regulation mechanism for **pp**MAPK^Spk1^ is robust and resistant to Ras1.G17V. It was also noted that the peak intensity of the **pp**MAPK^Spk1^ was somewhat lower and the rate of signal reduction was slightly decreased compared to wildtype cells (Fig. 1G).

### (4) The elongated *ras1.G17V* shmoos develop with a minimum level of MAPK^Spk1^: neither amplitude nor duration of ppMAPK^Spk1^ signal influences the *ras1.G17V* phenotype

Having observed that *MAPKK^byr1.DD^* and *ras1.G17V* show different MAPK^Spk1^ activation profiles (sustained vs transient) and different morphological phenotypes (*“fus” vs* elongated), we examined a link between the MAPK^Spk1^ activation profiles and the cell morphology. The MAPK^Spk1^ phosphorylation profile of the cells harbouring both *ras1.G17V* and *MAPKK^byr1.DD^* mutations (*ras1.G17V MAPKK^byr1.DD^* double mutant) was comparable to the cells harbouring *MAPKK^byr1.DD^* mutation only; it showed a slow increase of the **pp**MAPK^Spk1^-GFP level, which reached the plateau at about 16 hours after induction of mating (Fig. 2A, red line, original membrane images in Fig. S3). The nuclear **pp**MAPK^Spk1^-GFP signal in the *ras1.G17V MAPKK^byr1.DD^* double mutant was also present 24 hours after induction of mating, unlike the *ras1.G17V* single mutant cells, confirming that the *ras1.G17V MAPKK^byr1.DD^* double mutant cells retained a high **pp**MAPK^Spk1^ level. Thus, in terms of the activation status of MAPK^Spk1^, *MAPKK^byr1.DD^* is epistatic to *ras1.G17V*.

**Figure 2.**
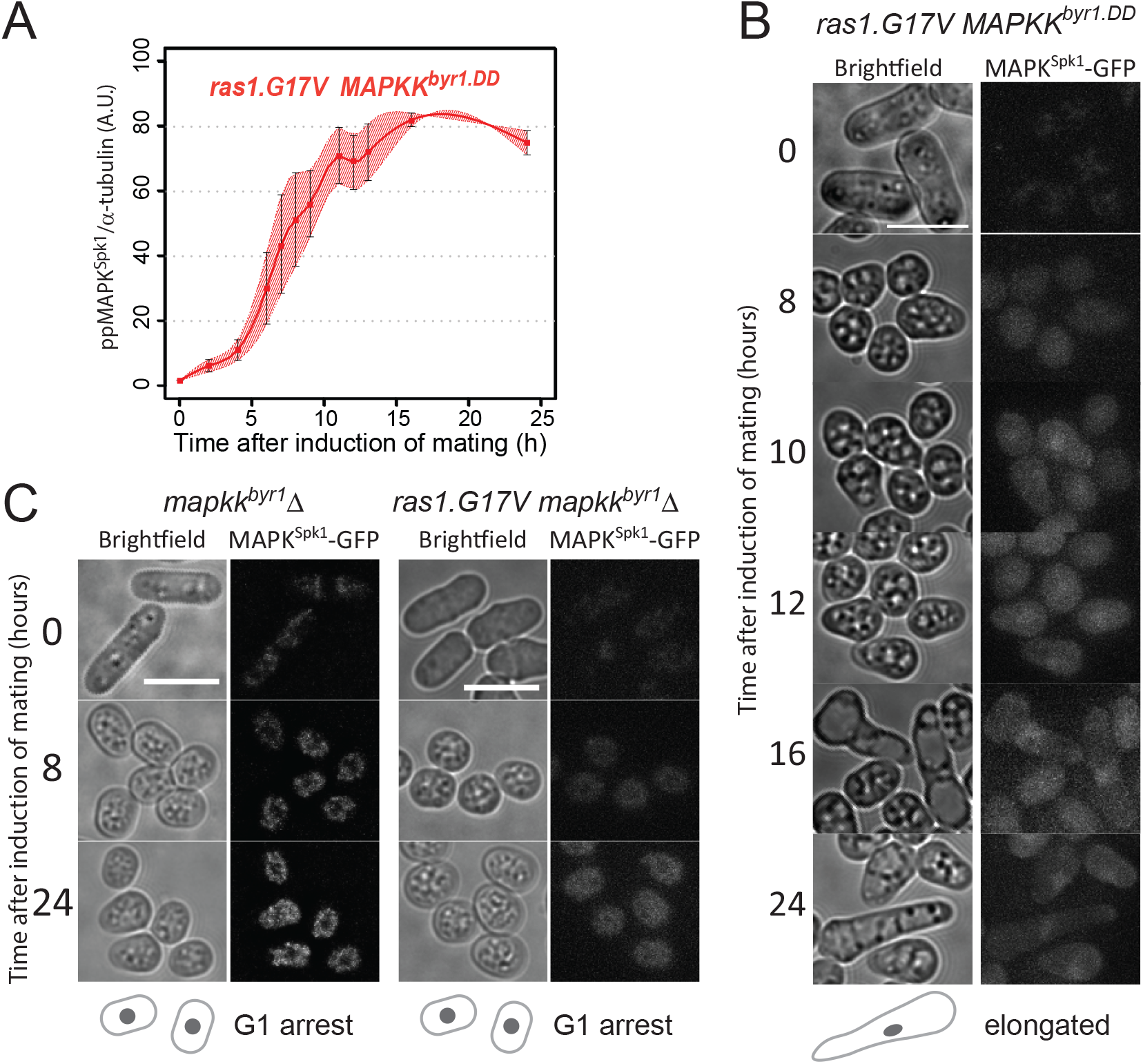
In the *ras1.G17V MAPKK^byr1.DD^* double mutant, the MAPK^Spk1^ phosphorylation profile follows *MAPKK^byr1.DD^* single mutant phenotype whilst cell morphology mimics the *ras1.G17V* single mutant phenotype. (A) MAPK^Spk1^ phosphorylation status in the *ras1.G17V MAPKK^byr1.DD^* double mutant cells (KT3439). Cells were induced for mating by the plate mating assay system as described in the materials and methods. Quantitated **pp**MAPK^Spk1^ signal (arbitrary unit) from western blots is presented. Results of two biological replicates (each derived from three technical replicates, error bars are ±SEM) are presented in red. (B) The terminal mating phenotype of *ras1.G17V MAPKK^byr1.DD^* double mutant is a phenocopy of *ras1.G17V* single mutant which shows the “elongated” morphology. Images were taken of *ras1.G17V MAPKK^byr1.DD^* double mutant (KT3439) in the same way as in Fig. 1. Time after induction of mating in hours is indicated on the left. (C) There is no morphological change in the absence of MAPK^Spk1^ signalling. Cell images of *MAPKK^byr1^Δ* (KT4700) and *ras1.G17V MAPKK^byr1^Δ* (KT5030) strains are shown. Images were taken in the same way as in Fig. 1. Time after induction of mating in hours is indicated on the left of each series. Scale bars represent 10μm.

In terms of cell morphology, the *ras1.G17V MAPKK^byr1.DD^* double mutant cells showed the “elongated” *ras1.G17V* phenotype (Fig. 2B), but not the “paired” *MAPKK^byr1.DD^* phenotype. Therefore, *ras1.G17V* was epistatic to *MAPKK^byr1.DD^* in terms of the elongated shmoo morphology.

Interestingly, obvious shmoo formation in the *ras1.G17V MAPKK^byr1.DD^* double mutant was first noticed 16 hours after induction of mating, much later than the *ras1.G17V* single mutant, but as a similar timing as the *MAPKK^byr1.DD^* single mutant (Fig. 2B). Given the slow increase of the **pp**MAPK^Spk1^ signal in mutants harbouring the *MAPKK^byr1.DD^* mutation, we predicted that the delayed appearance of the *ras1.G17V* shmoo meant that the *ras1.G17V* shmoo formation still requires a certain level of MAPK^Spk1^ activity. Indeed, when *MAPKK^byr1^* was deleted in the *ras1.G17V* mutant, not only was the MAPK^Spk1^ activation and nuclear MAPK^Spk1^-GFP signal abolished (Fig. S1F), but also shmoo formation was abrogated as in the *MAPKK^byr1^Δ* single mutant (Fig. 2C). Based on these observations, we concluded that the cell morphology is determined by the molecular status of Ras1, and not by the MAPK^Spk1^ activation profile. Yet, the *ras1.G17V* phenotype still requires MAPK^Spk1^ activity, which determines the timing of shmoo formation.

### (5) Cdc42 is required for the shmoo formation but not for the MAPK^Spk1^ activation

During the vegetative cycle, *ras1Δ* cells show spherical cell morphology (Fukui et al., 1986; Nadin-Davis et al., 1986a) and polarised localisation of active Cdc42 is compromised (Kelly and Nurse, 2011), (Fig. 4C, D), indicating that Ras1 is involved in Cdc42 activation. The GTP-loaded Cdc42 is then predicted to activate the downstream Ste20-like kinase, Pak1/Shk1 (Endo et al., 2003; Marcus et al., 1995; Ottilie et al., 1995; Verde et al., 1995), resulting in actin reorganisation and shmoo formation under mating conditions (Bendezu and Martin, 2013; Merlini et al., 2016).

To confirm that Cdc42 acts downstream of Ras1, we generated a double mutant strain harbouring *ras1.G17V* and deletion of *scd1*, encoding a GDP-GTP exchanging factor for Cdc42 (Cdc42-GEF^Scd1^). The *ras1.G17V* elongated shmoo phenotype was lost in the double mutant and instead, the cells showed a mating-deficient phenotype similar to the *cdc42-GEF^scd1^Δ* single mutant (Fig 3A). The result supports the model that Cdc42 acts downstream of Ras1 to cause morphological changes.

**Figure 3.**
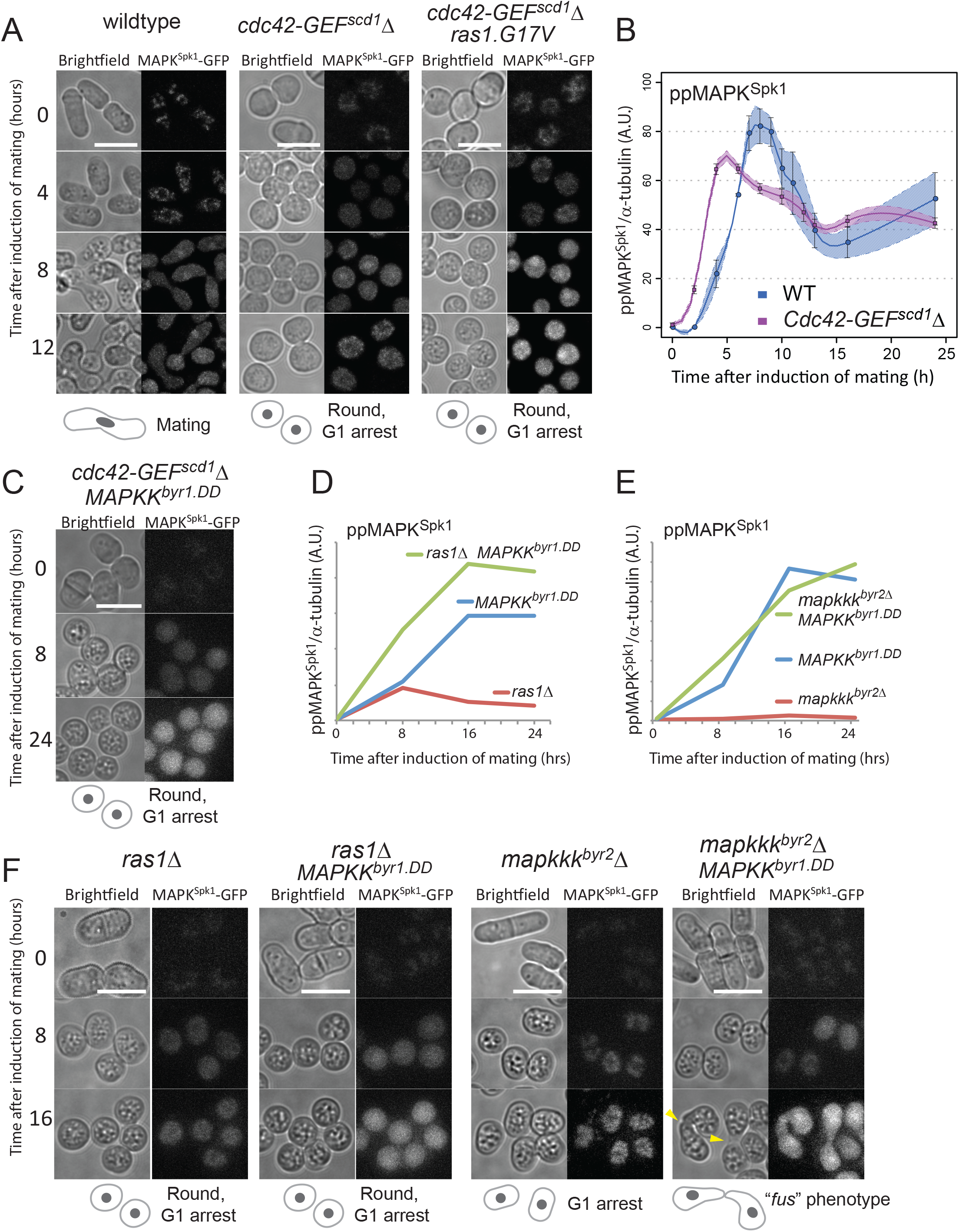
Ras1 activates both MAPK^Spk1^ and Cdc42 pathways during pheromone signalling. (A) *scd1Δ* morphology and MAPK^Spk1^-GFP signal. Images of WT (KT3082), *scd1Δ* (KT4061) and *scd1Δ ras1.G17V* double mutant (KT4056) were taken in the same way as in Fig.1. Numbers on the left represents hours after induction of mating. (B) MAPK^Spk1^ phosphorylation state in *scd1Δ* (KT4061) cells after mating induction. Results of three biological replicates (error bars are ±SEM) are presented. The wildtype **pp**MAPK^Spk1^ result presented in Fig.1 (C) is also shown in blue as a reference. (C) Cell images of *scd1Δ MAPKK^byr1.DD^* double mutant (KT4047) were taken in the same way as in Fig.1. Numbers on the left represents hours after induction of mating. (D) MAPK^Spk1^ phosphorylation state in *ras1Δ* (KT4323), *MAPKK^byr1.DD^* (KT3435) and *ras1Δ MAPKK^byr1.DD^* (KT4359) cell extracts. Original Western blotting data is presented in Fig. S4A. (E) MAPK^Spk1^ phosphorylation state in *mapkkk^byr2^Δ* (KT3763), *MAPKK^byr1.DD^* (KT3435) and *mapkkk^byr2^Δ MAPKK^byr1.DD^* (KT4010) cell extracts. Original Western blotting data is presented in Fig. S4B. For (D) and (E), quantification was carried out using the Image Studio ver2.1 (Li-cor). (F) Cell images of the strains mentioned in (D) and (E) were taken in the same way as in Fig.1. Numbers on the left represents hours after induction of mating. For all the images presented in (A), (C) and (F), scale bars represent 10μm.

Intriguingly, in the strains harbouring the *cdc42-GEF^scd1^Δ* mutation, a nuclear MAPK^Spk1^-GFP signal appeared (Fig. 3A), indicating that activation of MAPK^Spk1^ may not be impaired by lack of active Cdc42. This was unexpected because in a previous study, it was predicted that activation of Cdc42 contributes to activation of MAPKKK^Byr2^ (Tu et al., 1997). To clarify this issue, we measured **pp**MAPK^Spk1^ in the *cdc42-GEF^scd1^Δ* mutant. Strikingly, in these cells MAPK^Spk1^ activation occurred with a reproducible advancement of the initial activation timing compared to the wildtype cells (Fig. 3B, original membrane images in Fig. S3). The result shows that MAPK^Spk1^ activation does not require Cdc42 activity. Additionally, the faster activation of MAPK^Spk1^ raises the interesting possibility that two Ras1 effectors, MAPKKK^Byr2^ and Cdc42-GEF^Scd1^, are competing with each other for activated Ras1, thus, lack of Cdc42-GEF^Scd1^ results in an advanced MAPK^Spk1^ activation (modelled in Fig. 7A).

Substantial MAPK^Spk1^ activation in the *cdc42-GEF^scd1^Δ* mutant means that the mating deficiency of this mutant is unlikely to be the result of the lack of MAPK^Spk1^ activation. Indeed, introduction of *MAPKK^byr1.DD^* to the *cdc42-GEF^scd1^Δ* mutant did not restore the mating deficient phenotype, even though nuclear MAPK^Spk1^-GFP highly accumulated (Fig. 3C). The result shows that active Cdc42 function is absolutely required for the mating process regardless of the MAPK^Spk1^ activation status. Taken together with the essential role of MAPK^Spk1^, we concluded that the mating pheromone signalling feeds into at least two pathways, MAPK^Spk1^ and Cdc42.

### (6) Ras1 activates two effector pathways, MAPK^Spk1^ and Cdc42

In order to further clarify the role of Ras1 we examined the MAPK^Spk1^ activation status and cell morphology in the following four strains: *ras1Δ* mutant, *ras1Δ MAPKK^byr1.DD^* double mutant, *MAPKKK^byr2^Δ* mutant and *MAPKKK^byr2^Δ MAPKK^byr1.DD^* double mutant. As mentioned earlier, deletion of *ras1* causes cells to show a round morphology (Fig. 3F), with reduced cortical signal of CRIB-GFP, an indicator of the active GTP-bound form of Cdc42, showing that Cdc42 activation is compromised (Fig.4C, D). *ras1* deletion also causes substantial reduction, but not complete elimination, of the **pp**MAPK^Spk1^ (Fig. 3D, red line, and Supplementary Fig. S4A); thus, Ras1 plays an important role in activating both Cdc42 and MAPK^Spk1^ pathways. Introduction of the *MAPKK^byr1.DD^* mutation into the *ras1Δ* mutant cells induces the constitutive **pp**MAPK^Spk1^ (Fig. 3D, green line and Supplementary Fig. S4A) but does not affect the round cell morphology and cells remain sterile (Fig. 3F, the 2^nd^ left panel. Note the accumulating MAPK^Spk1^-GFP at 16 hours after induction of mating), as was the case for the *Cdc42-GEF^Scd1^Δ MAPKK^byr1.DD^* double mutant (Fig. 3C).

**Figure 4.**
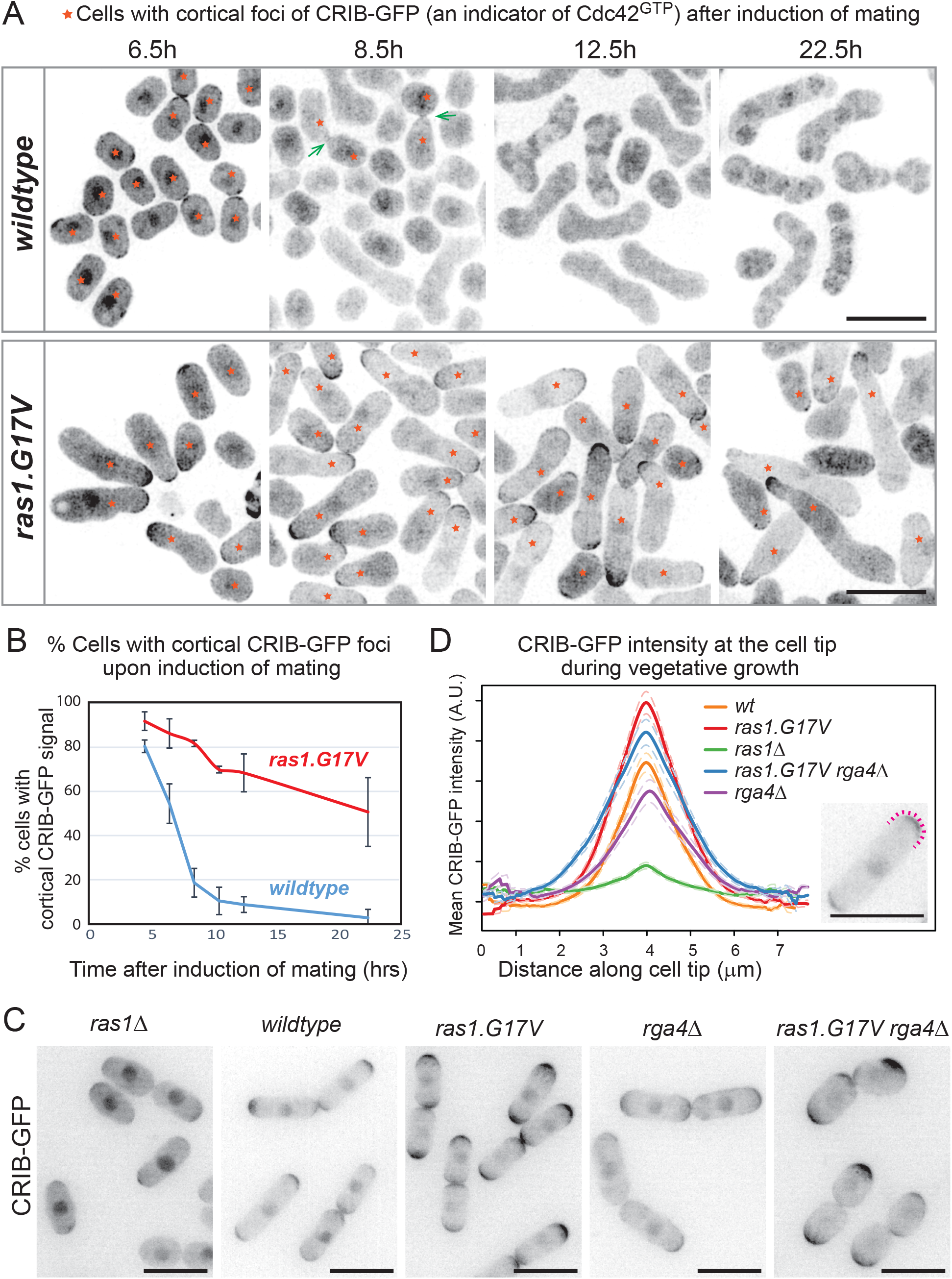
Ras1.G17V induces cortical Cdc42^GTP^ accumulation. (A) Cell morphology and localisation of Cdc42^GTP^, indicated by CRIB-GFP signal, during the sexual differentiation process. Wildtype (KT5077) and *ras1.G17V* (KT5082) mutant cells were induced for mating/sexual differentiation by the plate mating assay condition (Materials and Methods) and live cell images were taken at the indicated time after induction of mating/sexual differentiation. Representative CRIB-GFP signal images are presented. Cells with cortical CRIB-GFP foci are indicated by orange stars. Rapidly-disappearing CRIB-GFP signals at the fusion site of wildtype mating cells are indicated by green arrows at time 8.5h image. Scale bar: 10 μm. (B) Quantitation of the results presented in (A). At each time point (4.5h, 6.5h, 10.5h, 12.5h and 22.5h after induction of mating/sexual differentiation), 150 cells were examined whether they have cortical CRIB-GFP foci. % cells with cortical CRIB-GFP foci is presented. The experiment was repeated for three times and the mean values and SDs are plotted in the graph. (C) Cell morphology and localisation of Cdc42^GTP^, indicated by CRIB-GFP signal, during vegetative growth. Representative CRIB-GFP signal images of cells of wildtype (KT5077), *ras1Δ* (5107), *ras1.G17V*(KT5082), *rga4Δ* (5551) and *rga4Δ ras1.G17V*(KT5554) are presented. Scale bar: 10 μm. (D) Quantitated CRIB-GFP signals on the cell cortex of cells presented in (C). Intensity of GFP signal on the cell cortex was measured along one of the cell tips as indicated as a magenta dotted line in the example image on the right (Scale bar: 10 μm) as stated in the Materials and Methods. 40 cells without septum were measured for each strain and the average curve from all aligned traces per strain was calculated, and displayed with respective standard error of the mean curves (dashed lines) as described in Materials and Methods.

In a striking contrast, the sterile phenotype of the *MAPKKK^byr2^Δ*, associated with complete lack of shmoo formation (Fig. 3F, the 2^nd^ right panel), was converted to the *“fus”* phenotype, when combined with the *MAPKK^byr1.DD^* mutation (Fig. 3F, the far right panel). As expected, the *MAPKKK^byr2^Δ MAPKK^byr1.DD^* double mutant shows MAPK^Spk1^ constitutive activation (Fig. 3E and Supplementary Fig. S4B). Thus, unlike the cases of *scd1Δ* or *ras1Δ*, lack of *MAPKKK^byr2^* can be bypassed by constitutive activation of MAPK^Spk1^, indicating that the sole role of MAPKKK^Byr2^ is to activate the MAPK^Spk1^ unlike its upstream activator, Ras1, which also activates Cdc42 pathway (a model presented in Fig. 7A).

### (7) Ras1.G17V causes accumulation of Cdc42-GTP at the cell cortex

Having observed a relatively mild influence of Ras1.G17V towards the MAPK^Spk1^ activation, we next examined whether the Cdc42 pathway was affected by the *ras1.G17V* mutation. We visualized the active GTP-bound form of Cdc42 (Cdc42^GTP^) using CRIB-GFP that specifically binds to Cdc42^GTP^ (Tatebe et al., 2008). As previously observed, dynamic foci of CRIB-GFP appeared on the cell cortex upon induction of mating (Bendezu and Martin, 2013)(Fig. 4A and B). In our experimental condition, more than 80% of wildtype cells showed the cortical CRIB-GFP signal at 4.5 hours after induction of mating (Fig. 4B). The cortical CRIB-GFP foci became concentrated at the site of mating and quickly disappeared once cells fused successfully to form zygotes (Fig. 4A and B). In striking contrast, in the *ras1.G17V* mutant cells, the cortical CRIB-GFP signal persisted, often at the elongated tip end of the cells, even 12.5 hours after induction of mating (Fig. 4A and B). The signal could still be seen in about 40% of the cells 22.5 hours after induction of mating (Fig. 4A and B). The result shows that the Cdc42 pathway is excessively activated in the *ras1.G17V* mutant and the localisation pattern of Cdc42^GTP^ indicates that the signature “elongated” *ras1.G17V* morphological phenotype is caused by deregulation of the Cdc42 pathway.

Ras1-mediated Cdc42 pathway activation has been also indicated during the vegetative growth where the *ras1Δ* mutant shows a round cell morphology (Chang et al., 1994; Kelly and Nurse, 2011). However, unlike during the mating process, the *ras1.G17V* mutation does not cause an obvious morphological phenotype during the vegetative growth. We predicted that, during the vegetative growth, rigorous negative regulation occurs for Cdc42 by GTPase activation protein(s) (GAPs), such as Rga4 (Das et al., 2007; Kelly and Nurse, 2011; Tatebe et al., 2008) to counteract the effect of *ras1.G17V*. To examine this possibility, we compared Cdc42 activation status of vegetatively growing wildtype, *ras1Δ, ras1.G17V, rga4Δ*, and *rga4Δ ras1.G17V* double mutant cells (Fig. 4C and D). As previously described, CRIB-GFP showed a clearly polarized signal at the growing cell tips in the wildtype strain (Tatebe et al., 2008)(Fig. 4C and D). In the *ras1Δ* mutant, the cells were round and CRIB-GFP signal on the cell cortex had largely disappeared as was seen in the *cdc42-GEF^scd1^Δ* mutant (Kelly and Nurse, 2011). In contrast, the cortical CRIB-GFP signal was clearly increased in the *ras1.G17V* single mutant although the cell morphology appeared largely similar to the wildtype cells (Fig. 4C, D). These results indicate a direct involvement of Ras1 in activating Cdc42. Meanwhile, the *rga4Δ* mutant cells showed slight alterations to the cell morphology, accompanied with less polarized distribution of cortical CRIB-GFP signal, as has been reported (Fig. 4C and D) (Das et al., 2007; Kelly and Nurse, 2011; Tatebe et al., 2008). Strikingly, the *rga4Δ ras1.G17V* double mutant showed a clear morphological alteration (big round cells) and the strongest cortical CRIB-GFP signal among all the mutants examined (Fig. 4C and D). The result fits well with our hypothesis that Ras1.G17V is activating Cdc42 even during vegetative growth, but the overall effect of Ras1.G17V is counteracted by the Cdc42-GAP, Rga4.

### (8) Ras1 and an adaptor protein Ste4 are both necessary to fully activate MAPKKK^Byr2^

Although Ras1 clearly plays the major role to activate MAPK^Spk1^, a marginal, but detectable level of **pp**MAPK^Spk1^ was still induced in the *ras1Δ* mutant (Fig. 3D and Supplementary Fig. S4A), indicating that there is a Ras1-independent mechanism to activate MAPK^Spk1^. Previous studies proposed an adaptor protein, Ste4, to be involved in the activation of MAPKKK^Byr2^(Barr et al., 1996; Okazaki et al., 1991; Ramachander et al., 2002; Tu et al., 1997). We therefore examined whether Ste4 is required for MAPK^Spk1^ activation.

In constrast to the *ras1Δ* mutant, we detected virtually no MAPK^Spk1^ phosphorylation in the *ste4Δ* mutant (Fig. 5A and Supplementary Figure S4C), indicating that the adaptor^Ste4^ is a prerequisite for the MAPK^Spk1^ activation and the **pp**MAPK^Spk1^ signal observed in the *ras1Δ* mutant is dependent on Ste4 function. Introduction of *ras1.G17V* mutation neither restored the MAPK^Spk1^ activation nor mating (Fig. 5A, B), thus, activation of Ras1 cannot take over Ste4 function. In a striking contrast, the *ste4Δ MAPKK^byr1.DD^* double mutant showed the *“fus”* phenotype as the *MAPKK^byr1.DD^* single mutant cells and induced constitutive MAPK^Spk1^ activation, indicating that Ste4 is solely required for MAPK^Spk1^ activation (Fig. 5A, B). Taken together, MAPKKK^Byr2^ is activated through a mechanism involving both Ras1 and Ste4, but Ste4 only conveys the signal towards the MAPK^Spk1^, while Ras1 also activates Cdc42.

**Fig. 5.**
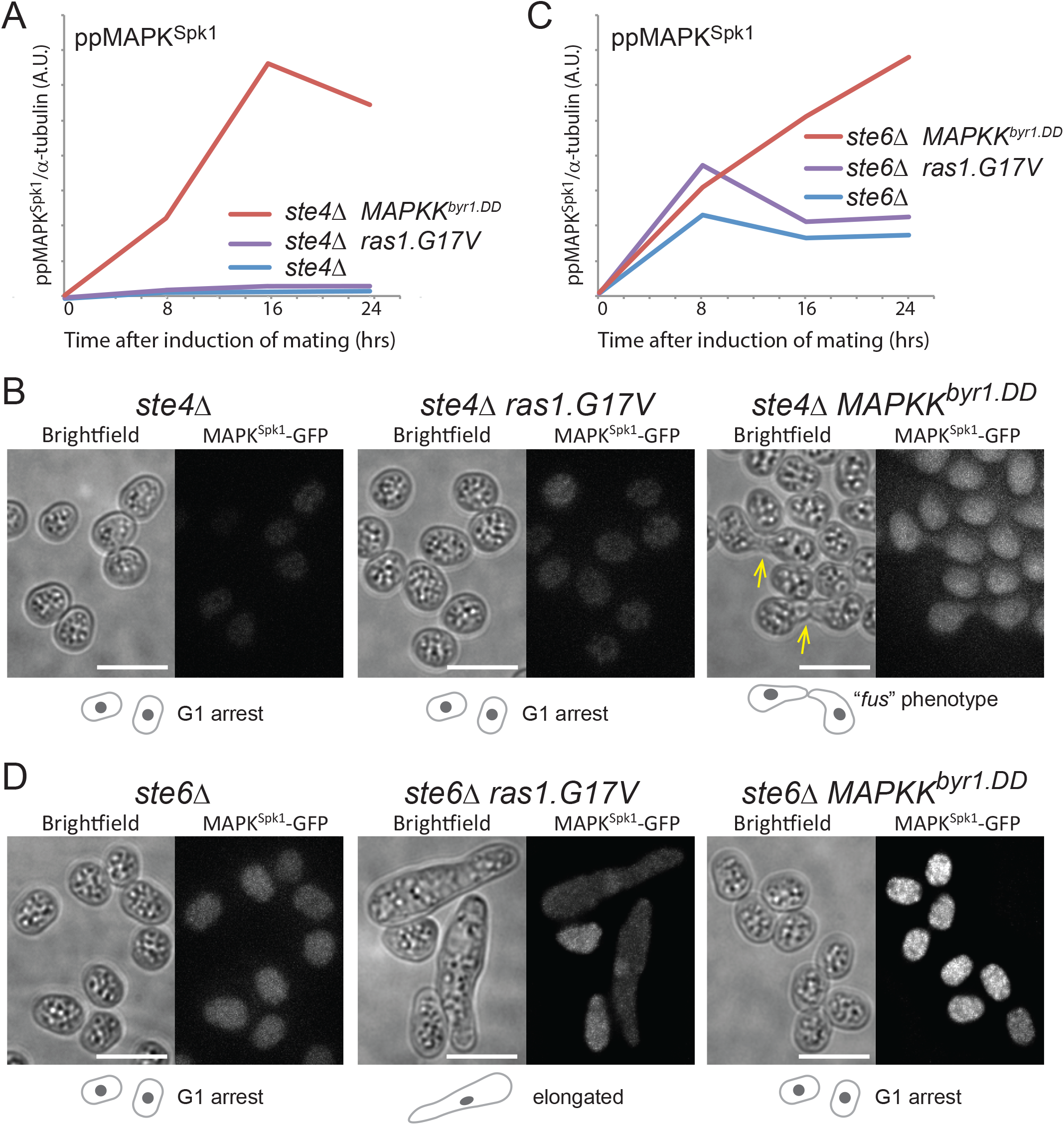
Distinct contributions of Ste4 and Ste6 to MAPK^Spk1^ phosphorylation. (A) Ste4 is essential for MAPK^Spk1^ activation. MAPK^Spk1^ phosphorylation status in *ste4Δ* (KT4376), *ste4Δ ras1.G17V*(KT5143) and *ste4Δ MAPKK^byr1.DD^* (KT5136) at times-points 0, 8, 16 and 24 hours post mating induction are presented. Original Western blotting membranes are presented in Fig. S4C. (B) Incapability of *ste4Δ* to cause pheromone-induced morphological change is suppressed by *MAPKK^byr1.DD^* but not by *ras1.G17V*. Cell images of *ste4Δ* (KT4376), *ste4Δ ras1.G17V* (KT5143) and *ste4Δ MAPKK^byr1.DD^* (KT5136) strains were taken 24 hours after induction of mating. Scale bar: 10 μm. (C) Lack of Ste6 does not result in the complete loss of MAPK^Spk1^ phosphorylation. MAPK^Spk1^ phosphorylation status in *ste6Δ* (KT4333), *ste6Δ ras1.G17V* (KT4998) and *ste6Δ MAPKK^byr1.DD^* (KT5139) at times-points 0, 8, 16 and 24 hours post mating induction are presented. Original Western blotting membranes are presented in Fig. S4D. (D) Incapability of *ste6Δ* to cause pheromone-induced morphological change is suppressed by *ras1.G17V* mutation but not by *MAPKK^byr1.DD^* mutation. Cell images of *ste6Δ* (KT4333), *ste6Δ ras1.G17V* (KT4998) and *ste6Δ MAPKK^byr1.DD^* (KT5139) strains were taken 24 hours after induction of mating. Scale bar: 10 μm.

### (9) Ste6, a Ras1 GTP-GDP exchange factor, contributes to both the MAPK^Spk1^ and the Cdc42 pathway activation

There are two GDP-GTP exchange factors (GEFs) identified for Ras1: Ste6 and Efc25 (Hughes et al., 1990; Tratner et al., 1997). As to the functional differences, Ste6 is essential for mating but is dispensable during the vegetative cycle whilst Efc25 is dispensable for mating but is required to maintain the cell morphology during the vegetative growth (Hughes et al., 1990; Tratner et al., 1997). There has been an interesting proposition that Ste6 may specifically help Ras1 to activate the MAPK^Spk1^ pathway, but not the Cdc42 pathway, whilst Efc25 specifically facilitates Ras1 to activate the Cdc42 pathway (Papadaki et al., 2002). We examined this hypothesis by monitoring the MAPK^Spk1^ activation status and conducting genetic epistasis analysis of *ras1.G17V* and *MAPKK^byr1.DD^* in the *ste6Δ* mutant.

In *ste6Δ* cells, MAPK^Spk1^ phosphorylation was found somewhat reduced but occurred at a clearly detectable level. The signal increased when the *ras1.G17V* mutation was introduced (Fig. 5C and Supplementary Figure S4D). When *ste6Δ and MAPKK^byr1.DD^* were combined, MAPK^Spk1^ signalling recapitulated the *MAPKK^byr1.DD^* activation profile (Fig. 5C and Supplementary Figure S4D). Nonetheless, the “pheromone-insensitive sterile” morphology of *ste6Δ* was only rescued by *ras1.G17V*, as previously reported, exhibiting the “elongated” phenotype (Hughes et al., 1990), but not by MAPKK^byr1.DD^ (Fig. 5D). The result indicates that, unlike *ste4Δ* mutant, the mating deficiency of *ste6Δ* is not caused by mere lack of MAPK^Spk1^ activation but by lack of Ras1 activation. We concluded that Ste6 functions to activate Ras1, which then activates *both* the MAPK and Cdc42 pathways in response to pheromone signalling.

### (10) Activation mutant of Gpa1 mimics the full pheromone signalling

In order to generate an integrated prototype Ras signalling model, we further investigated the upstream signal input machinery. Previous studies showed that Gpa1 plays the primary role in pheromone signalling (Obara et al., 1991). In agreement, in the *gpa1Δ* mutant we detected no MAPK^Spk1^ activation nor nuclear accumulation of MAPK^Spk1^-GFP (Fig. 6A, B and Supplementary Fig. S5A). Introducing the *ras1.G17V* to the *gpa1Δ* strain did not rescue the complete lack of *MAPK^Spk1^* activation nor did it induce a shmoo-like morphological change (Fig. 6A, B and Supplementary Fig. S5A), supporting our earlier observation that a Ras1-independent mechanism, involving Ste4 is essential for MAPK^Spk1^ activation. Meanwhile, introducing the *MAPKK^byr1.DD^* mutation caused the constitutive activation of MAPK^Spk1^ (Fig. 6A and Supplementary Fig. S5A), but cells showed no morphological change (Fig. 6B). When both *ras1.G17V* and *MAPKK^byr1.DD^* mutations were introduced into the *gpa1Δ* strain, MAPK^Spk1^ was activated and a shmoo-like morphological change occurred (Fig. 6A, B and Supplementary Fig. S5A). Therefore, activation of both of these two molecules is required and sufficient to mimic the pheromone signalling.

**Fig. 6.**
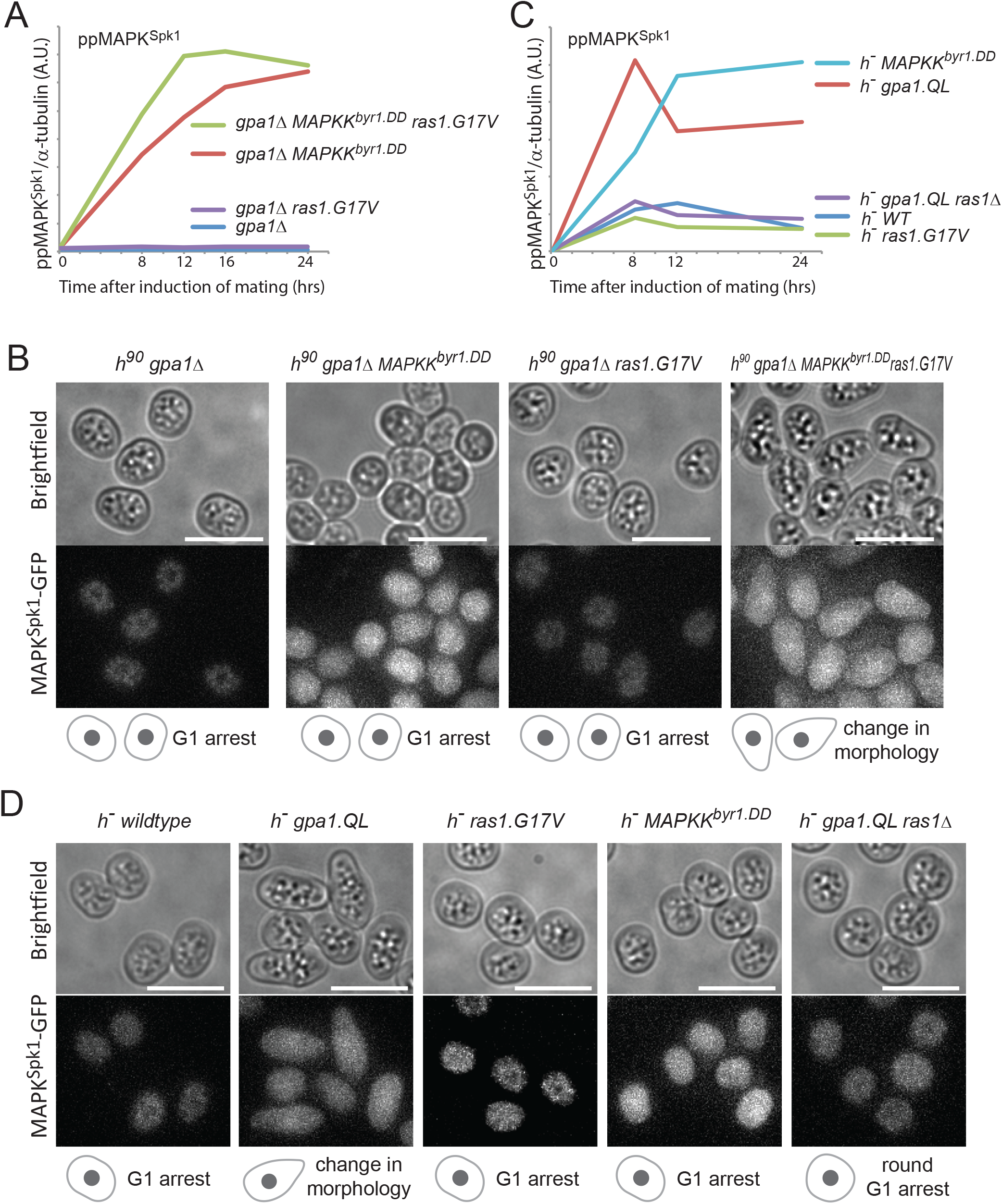
Gpa1 transduces the pheromone signalling by activating MAPK^Spk1^ and Ras1 pathways. (A) MAPK^Spk1^ phosphorylation status in homothallic *gpa1Δ* (KT4335), *gpa1Δ ras1.G17V* (KT5023), *gpa1Δ MAPKK^byr1.DD^* (KT4353) and *gpa1Δ ras1.val17 MAPKK^byr1.DD^* (KT5035) at times-points 0, 8, 12, 16 and 24 hours after mating induction. Original Western membrane is presented in Fig. S5A. (B) Cell images of the above mentioned strains at 16 hours after mating induction. All the cell images were taken and processed as in Figure 1. Scale bar is 10 μm. (C) MAPK^Spk1^ phosphorylation status in *h^−^* WT (KT4190), *h^−^ gpa1.QL* (KT5059), *h^−^ ras1.G17V* (KT4233), *h^−^ gpa1.QL ras1Δ* (KT5070) and h^−^ *MAPKK^byr1.DD^* (KT4194) at times-points 0, 8, 12 and 24 after mating induction. Note that while the activation induced in the *gpa1.QL* mutant was down-regulated, *MAPKK^byr1.DD^* induced a constitutive activation. Original Western membrane is presented in Fig. S5B. (D) Cell images of the above strains at 12 h after induction of mating. All the cell images were taken and processed as in Figure 1. Scale bar is 10 μm.

To further confirm that the Gpa1 is a central component of pheromone signalling, we looked into the MAPK^Spk1^ activation status in the constitutively active *gpa1.QL* mutant, which exhibits a “shmoo-like” morphological change in the heterothallic *h^−^* strain without the mating partner (Obara et al., 1991).

Upon nitrogen starvation, the *h^−^ gpa1.QL* mutant strain showed morphological changes and a strong MAPK^Spk1^ activation (Fig. 6C red line, Fig. 6D and Supplementary Fig. S5B). This response was largely dependent on the Ras1 function as the *h^−^ gpa1.QL ras1Δ* double mutant exhibited a significantly reduced level of **pp**MAPK^Spk1^ and a round cell morphology, two typical features of *ras1Δ* cells (Fig. 6C, D and Supplementary Fig. S5B). On the other hand, the *h^−^ ras1.G17V* single mutant showed a very low level of MAPK^Spk1^ activation, comparable to the one observed in the *h^−^* wildtype strain, with no apparent morphological alternation, confirming that sole activation of Ras1 does not substitute the pheromone signalling.

The *h^−^ MAPKK^byr1.DD^* mutant induced a strong constitutive MAPK^Spk1^ activation, confirming that the MAPKK^Byr1.DD^ molecule can activate MAPK^Spk1^ regardless of the pheromone signal input (Fig. 6C, light-blue line and Supplementary Fig. S5B). However, cell morphology was unchanged (Fig. 6D). Collectively, these results support the model where Gpa1 acts as the central transducer of the pheromone signalling, which can be mimicked only if both MAPK^Spk1^ and Ras1 are activated.

### (11) A holistic modelling framwork of MAPK^Spk1^ activation

Based on quantitative **pp**MAPK^Spk1^ measurements in wildtype and various mutant strains (Fig. 1C, E, G, and Fig. 3B), we constructed a mathematical model of the MAPK^Spk1^ signalling dynamics. The aim of the model is to test whether a simple competition of the MAPK and the Cdc42 pathways for a shared pool of active Ras^GTP^, can explain why the *scd1Δ* strain shows the similar **pp**MAPK^Spk1^ activation profile as the *ras1.G17V* strain.

We designed a reductionist model of 6 ordinary differential equations to represent key steps of pheromone signalling (Fig. 7A, Supplementary Fig. S6 and materials and methods). Model simulations and parameter estimations were performed in COPASI (Hoops et al., 2006) and details of the modelling process are described in Materials and Methods. Each biochemical process is referred as [L1]-[L10] as depicted in Fig. 7A. All the signalling components were set to be in close proximity, based on our own observation (Fig. 1 and Fig. 4) and recent series of localization studies (Dudin et al., 2016; Merlini et al., 2016; Merlini et al., 2018). This allowed the model to predict some signaling to involve protein-protein interactions.

**Fig. 7.**
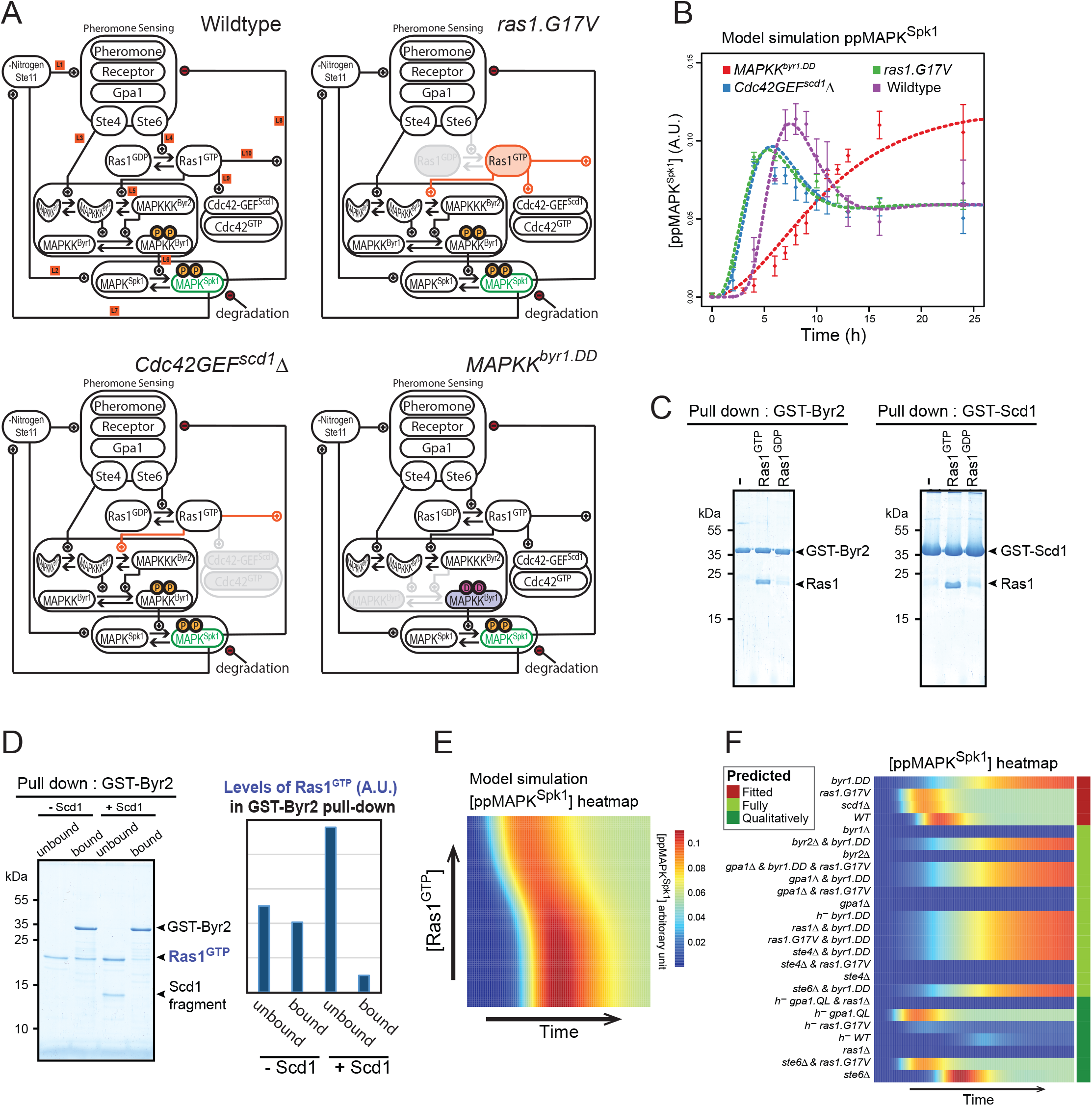
Mathematical modelling of the fission yeast pheromone pathway dynamics. (A) Components and frameworks of the mathematical model in wildtype and signalling mutants: *ras1.G17V, Cdc42GEF^scd1^Δ*, and *MAPKK^byr1.DD^*. Changes corresponding to each mutant are indicated as follows: Grey: removed components or interactions, orange: increased level of activity. For the exact implementation of the mutants, see Materials and Method. The measured component, **pp**MAPK^Spk1^, is highlighted in green. (B) Measured and simulated **pp**MAPK^Spk1^ activation profiles in wildtype, *ras1.G17V, Cdc42GEF^scd1^Δ* and *MAPKK^byr1.DD^* mutants. Dashed lines: model simulations. Diamonds: experimental data presented in Fig. 1C, E, G, and Fig. 3B; error bars: SEM. (C) GTP-loaded Ras1.G17V (1-172) directly binds to Byr2 (65-180) and Scd1 (760-872). *In vitro* GST pull-down assays of bacterially expressed Ras1.G17V (1-172), GST-Byr2 (65-180) and GST-Scd1 (760-872) were conducted as described in materials and methods. GTP-loaded Ras1.G17V (1-172) was found to bind to both GST-Byr2 (65-180) and GST-Scd1 (760-872). (D) Two Ras1 effectors, Byr2 and Scd1, compete for GTP-loaded Ras1.G17V (1-172). In vitro GST pull-down assays of bacterially expressed Ras1.G17V (1-172) and GST-Byr2 (65-180) were conducted as in (A). Addition of Scd1 (760-872) fragment interfered with Ras1-Byr2 binding (the 4^th^ lane). Quantitated signal intensities of the Ras1.G17V (1-172) band in the gel are shown in the right panel. (E) Simulated ppMAPK^Spk1^ dynamics in the wildtype model at increasing concentrations of Ras1^GTP^ added *in silico* to the system. Increased Ras1^GTP^ concentration causes advanced and reduced **pp**MAPK^Spk1^ peak intensities. (F) The model fitted to the 4 strains (as above, in red) correctly predicts ppMAPK^Spk1^ dynamics in the additional 21 signalling mutant strains (in light and dark green) measured in this study.

The framework of the modelling process is as follows: Genes encoding pheromones, receptors, Gpa1, Ste4 and Ste6 are all known to be under regulation of Ste11, the master transcriptional regulator for meiotic genes (Hughes et al., 1994; Mata and Bahler, 2006; Mata et al., 2002; Mata et al., 2007; Sugimoto et al., 1991) and these components are grouped into the Pheromone Sensing (PS) unit. During the vegetative growth, the PS unit is set to zero. Nitrogen starvation activates Ste11 (Kjaerulff et al., 2007; Sugimoto et al., 1991), which induces the PS unit [L1]. The PS unit activates MAPKKK^Byr2^ in a twofold manner: Directly by Ste4 [L3] and through Ras1 [L4](Fig. 7A). Activated MAPKKK^Byr2^ then triggers activation of MAPKK^Byr1^ [L5] that activates MAPK^Spk1^ [L6]. Since activated MAPK^Spk1^ further activates Ste11 (Kjaerulff et al., 2005), MAPK^Spk1^ has a positive feedback loop on its own expression via Ste11 [L2,L7] (Fig. 7A). As the pheromone signalling was found to induce a transient **pp**MAPK^Spk1^ peak (Fig. 1C), **pp**MAPK^Spk1^ activity is ought to be regulated by a delayed downregulation. Because the *MAPKK^byr1.DD^* mutant completely lacks downregulation (Fig. 1E), downregulation occurring downstream of MAPKK^byr1^ (e.g.: Pyp1 and Pmp1) were considered physiologically insignificant. Meanwhile, Sxa2 (a serine carboxypeptidase against a mating pheromone P-factor) and Rgs1 (a regulator of Gpa1), both of which are induced upon successful pheromone signalling (Imai and Yamamoto, 1992; Mata and Bahler, 2006; Pereira and Jones, 2001; Watson et al., 1999), receptor internalization (Hirota et al., 2001) and regulation of the *mapk^spk1^* transcript or other components by antisense RNA (Bitton et al., 2011) fit well to the criteria for the negative feedback. We represented all these potential negative feedbacks collectively as a single circuit, [L8]. Importantly, this downregulation [L8] works unperturbed in the presence of Ras1.G17V, by acting through the Ste4-dependent MAPKKK^Byr2^ activation process (Fig. 1G, Fig. 7A). Ras1-GTP activates both the MAPKKK^Byr2^ and the Cdc42 pathways [L9] (Chang et al., 1994). As opposed to previous expectations that active Cdc42^GTP^ is required to activate MAPKKK^Byr2^ (Tu et al., 1997), deletion of Cdc42-GEF^Scd1^ does not compromise MAPK^Spk1^ activation (Fig. 3B) and rather, it makes **pp**MAPK^Spk1^ dynamics remarkably similar to that of the Ras1.G17V strain (Fig. 3B and Fig. 1G, plotted together in Fig. 7B): in both cases, **pp**MAPK^Spk1^ peaks earlier than the wildtype case. At a molecular level, reactions depleting the Ras1^GTP^ pool are compromised in both strains. Therefore, to explain these observations, we hypothesized that MAPK^Spk1^ and Cdc42 pathways are competing for the common Ras1^GTP^ pool. In this manner, one of the pathways can modulate the other by changing the amount of unbound (available) Ras1^GTP^. In support of this prediction, we showed that binding of recombinant Ras1.G17V^GTP^ to a MAPKKK^Byr2^ fragment was reduced by the presence of a Cdc42-GEF^Scd1^ fragment *in vitro* (Fig. 7C, D). In this assay, bacterially expressed GST tagged fragments of both MAPKKK^Byr2^ (65-180) and Cdc42-GEF^Scd1^ (760-872) showed a specific binding towards the GTP-loaded Ras1.G17V (1-172) (Fig. 7C). However, the binding of Ras1.G17V^GTP^ to MAPKKK^Byr2^ was substantially decreased when the Cdc42-GEF^Scd1^ fragment was added (Fig. 7D). The result likely refects the intrinsic biochemical competitive nature of MAPKKK^Byr2^ and Cdc42-GEF^Scd1^ for Ras1 binding. As we show below, this simple hypothesis successfully describes pheromone signalling mutants tested in this study, suggesting that no unproven cross links are necessary to reproduce the *in vivo* observations.

An intriguing common feature observed in both Cdc42-GEF^scd1^Δ and *ras1.G17V* mutants is that **pp**MAPK^Spk1^ peaks not only at an earlier time point, but also with a *lower amplitude* as compared to the wildtype. If increased Ras1^GTP^ levels simply accelerate MAPK^Spk1^ activation, as has been conventionally assumed, **pp**MAPK^Spk1^ production should peak earlier *and higher* (Supplementary Fig S7B, best fit out of 1000 global fits), and the addition of Ras1^GTP^ only should increase the amplitude, but not affect timing (Supplementary Fig S7C). We confirmed these results in the best models from 1000 global fits (Materials and Methods). The comparison between our experimental results and the *in silico* predictions suggests that the role of Ras1^GTP^ is more complex than previously thought.

Strikingly, if we hypothesize that Ras1^GTP^ also contributes to the negative feedback [L10] (Fig. 7A), we recapitulate the *“earlier and lower”* peak of **pp**MAPK^Spk1^ in both the *ras1.G17V* and the Cdc42-GEF^scd1^Δ mutants (Fig. 7B). We currently do not have a direct experimental evidence to support [L10]. However, considering the fact that Ras1^GTP^ likely acts as a physical signalling hub at the cell cortex, mediating MAPKKK^Byr2^ recruitment and activation, which leads to recruitment of MAPKK^Byr1^ and MAPK^Spk1^, Ras1^GTP^ may work as a two-way amplifier, both assisting localized MAPK^Spk1^ activation at the shmoo site, as well as helping the negative feedback by concentrating the affected molecules.

The model successfully recapitulated the experimental results for wildtype, *ras1.G17V*, MAPKK^Byr1.DD^ and *cdc42-GEF^scd1^Δ* mutants (Fig. 7B). To test the predictive capacity of the model, we next performed an *in silico* experiment where we titrated increasing amounts of Ras1^GTP^ in the wildtype condition before nitorgen removal. In agreement with our hypothesis, we obtained a *ras1.G17V-like* **pp**MAPK^Spk1^ activation profile with increasing amount of Ras1^GTP^, i.e., the **pp**MAPK^Spk1^ peaks earlier with a lower peak intensity (Fig. 7E). The result further supported that Ras1^GTP^ availability alone is sufficient to explain both the *ras1.G17V* and *Cdc42-GEF^scd1^Δ* phenotypes.

To further test the predictive value of the model, we asked whether it could predict **pp**MAPK^Spk1^ dynamics in the 21 other strains, which were measured (Fig. 2-6), but not used for fitting the model. We implemented each mutation in the wildtype model (Supplementary Table S2) and the model accurately predicted relative **pp**MAPK^Spk1^ dynamics in 17 cases, or showed predicitons in close proximity to the observed **pp**MAPK^Spk1^ dynamics in the 4 remaining cases (Fig. 7F). Concluding from these results, our model likely represents the physiological framework of fission yeast RAS-MAPK signalling.

## Discussion

By quantitating the MAPK^Spk1^ and Cdc42 activation status during the mating process and conducting epistasis analysis between numerous signalling mutants, we showed that Ras1 coordinates activation of both the MAPK^Spk1^ cascade and the Cdc42 pathway. Furthermore, we revealed that the *ras1.G17V* mutant phenotype is caused by deregulation of Cdc42, rather than altered activation of MAPK^Spk1^ in physiological setting. Based on the experimental data, we built a mathematical model, incorporating an assumption that two Ras1 effectors, Cdc42-GEF^Scd1^ and MAPKKK^Byr2^, are competing for active Ras1. This is the first simulation of a physiological Ras signalling network in a whole organism. The model faithfully recapitulates MAPK^Spk1^ activation profiles in the wildtype and all mutant strains examined in this study. The model implies that targeting one of the RAS effector pathways can potentially result in a complex outcome, rather than simply shutting down the targeted effector pathway. We concluded that fission yeast pheromone Ras signalling is not only defined by compartmentalisation (Onken et al., 2006) but rather a coordination of events involving both the MAPK^Spk1^ and Cdc42 pathways. Coordinated activation of MAPK and Cdc42 pathways upon pheromone stimulation is a common feature shared by budding yeast pheromone signalling. In budding yeast, however, Ras homologues are not involved in the process. β and γ subunits of the receptor-coupled trimeric G-protein act as the hub to activate both pathways ((Hegemann et al., 2015; McClure et al., 2015), reviewed in (Atay and Skotheim, 2017)). It is intriguing that the key hub moleucules are evolutionary diverged in these yeast species whilst the effector pathways are conserved to bring comparable biological outcomes.

In this study, we confirmed that Gpa1, the α subunit of the receptor-coupled trimeric G-protein, is the central player of the pheromone signalling. It is likely that Gpa1 is the most downstream molecule conveying the complete pheromone signal. Considering that all the pheromone signalling components examined so far have been found at the shmoo site (Dudin et al., 2016; Merlini et al., 2016; Merlini et al., 2018)(this study), an attractive hypothesis is as follows: firstly, the activated pheromone receptor Map3/Mam2 locally activates Gpa1, which activates Ras1 and MAPK^Spk1^. This then leads to a localised activation of the Cdc42, causing shmoo formation in the direction of a mating partner (Fig. 8A).

**Fig. 8.**
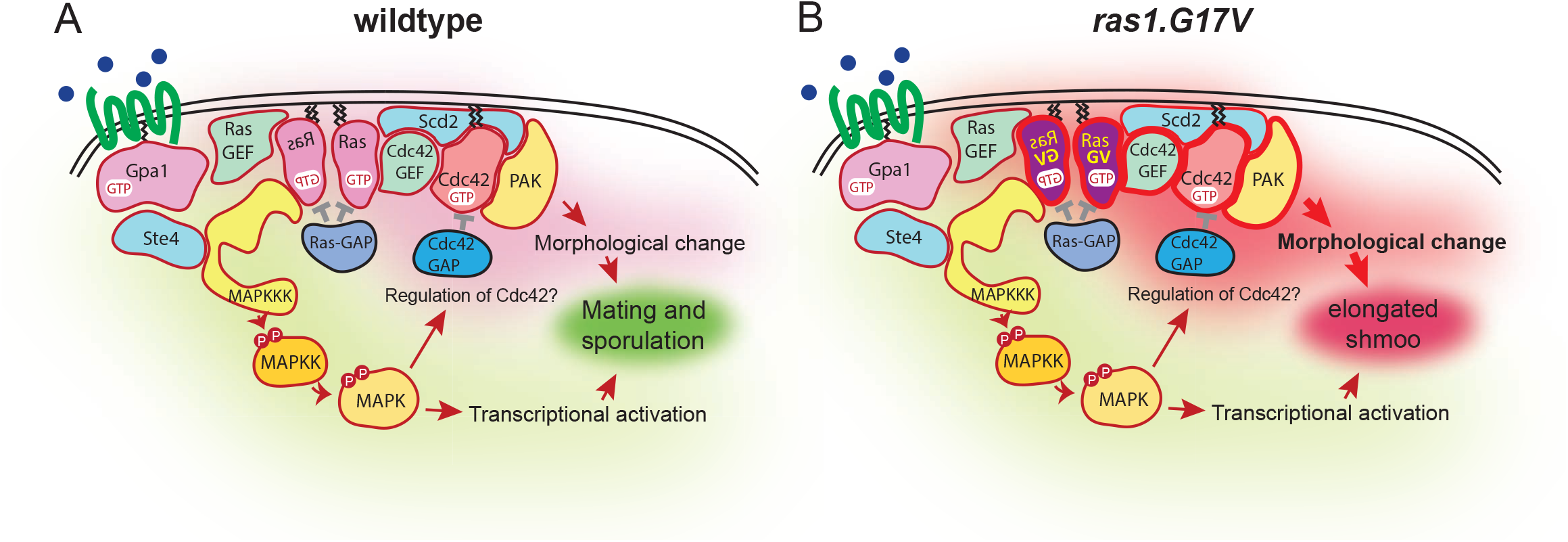
Schematic diagram of the fission yeast pheromone signalling pathway. (A) Wildtype fission yeast pheromone signalling pathway, highlighting the branched pheromone sensing, and that both branches are necessary for mating. (B) Because of the robust regulatory mechanism implemented for the MAPK pathway and the competition between the effectors, the constitutively active Ras1.G17V mutation yields a biased overactivation of Cdc42.

The *ras1.G17V* mutation led to an immediate activation of MAPK^Spk1^ compared with the wildtype cells. The G17V mutation of Ras1 is equivalent of G12V mutation of mammalian RAS, which results in a substantial reduction of both intrinsic and GAP-mediated GTPase activities (Trahey and McCormick, 1987). Therefore a larger fraction of Ras1 is expected to be in the active GTP-bound form in the *ras1.G17V* mutant cells. By mathematical modelling, we showed that the increased Ras1^GTP^ pool can explain a faster activation of MAPK^Spk1^. To our surprise, the constitutive Ras1.G17V mutation did not induce over-activation of MAPK^Spk1^. The attenuated MAPK^Spk1^ activation in the presence of Ras1.G17V indicated that an efficient feedback mechanism is in place to counteract the effect of Ras1.G17V. Strikingly, the same trend has been reported in the mouse model of the *K-ras^G12D^* mutation integrated at the endogenous chromosome locus (Tuveson et al., 2004). Therefore it is highly likely that the MAPK cascade is generally robust against upstream oncogenic constitutive stimulation. Based on our observation that the MAPK^Spk1^ is constitutively activated in the *MAPKK^byr1.DD^* mutant, we predict that the negative regulation occurs upstream of, or at the same level as, MAPKK^Byr1^, rather than phosphatases that directly regulate MAPK^Spk1^. In humans, ERK is shown to phosphorylate RAF proteins, the prototype MAPKKKs, to contribute to ERK signal attenuation (Brummer et al., 2003; Dougherty et al., 2005; Ritt et al., 2010). In future studies it will be important to determine whether MAPK^Spk1^ can directly downregulate MAPKKK^Byr2^ in a physiological setting.

Our results also show that an adaptor^Ste4^ plays a crucial role in activating MAPKKK^Byr2^, abolishing **pp**MAPK^Spk1^ production even in the presence of *ras1.G17V* mutation. This suggests that the adaptor^Ste4^ fits well to be one of the major targets by the negative feedback loop against **pp**MAPK^Spk1^. This mechanism is shared by budding yeast, where an adaptor protein, Ste50, modulates MAPKKK^Ste11^ (Ramezani-Rad, 2003). In humans, although such an adaptor protein for RAF proteins has yet to be identified, multiple RAF-interacting proteins, including 14-3-3 proteins, as well as formation of heterodimers between BRAF and CRAF, have been studied for their Ras-independent mechanism to activate RAF proteins (Lavoie and Therrien, 2015). Collectively, MAPK cascades seem to retain a general resistance to oncogenic RAS mutations in physiological settings (Fig. 8B).

Whilst lack of Cdc42 activation does not impair MAPK^Spk1^ activation, we found that MAPK^Spk1^ activity is required for shmoo formation even in the presence of Ras1.G17V (Fig. 2C). Therefore, the two Ras1 effectors, Cdc42 and MAPK^Spk1^ pathways, are not completely separable. The situation is reminiscent of the *K-ras^G12D^* MEFs (Tuveson et al., 2004). In this system, the *K-ras^G12D^* MEFs showed morphological anomalies. As both ERK and AKT phosphorylation levels in the *K-ras^G12D^* MEFs resembled wildtype, these pathways unlikely caused the morphological phenotype. Nonetheless, inhibitors against MAPK and PI3K pathways reverted the *K-ras^G12D^*–induced abnormal morphology back to the one similar to wildtype. The observation suggests that MAPK and PI3K pathways somehow contribute to the *K-ras^G12D^* morphological phenotype; for example, a basal level of MAPK and PI3K pathway activation may be a prerequisite for the *K-ras^G12D^*–induced morphological anomalies.

The molecular mechanism of how MAPK^Spk1^ contributes to Cdc42 activation will require further studies. Key components in pheromone signalling are transcriptionally up-regulated upon pheromone signalling (Xue-Franzen et al., 2006). This is driven by MAPK^Spk1^, which activates the master transcriptional regulator Ste11 (Kjaerulff et al., 2005; Mata and Bahler, 2006; Xue-Franzen et al., 2006). Therefore, the contribution of MAPK^Spk1^ to Cdc42 activation is expected to occur, at least partly, through transcriptional activation.

In addition, localisation of signalling components may be regulated by MAPK^Spk1^. In budding yeast, MAPK^Fus3^ brings Cdc24, the GEF for Cdc42, to the shmoo site by phosphorylating an adaptor protein Far1 (Hegemann et al., 2015), which otherwise sequesters Cdc24 into the nucleus (Nern and Arkowitz, 2000; Shimada et al., 2000). Fission yeast does not have an obvious Far1 homologue but MAPK^Spk1^ may also directly phosphorylate and activate Cdc42 and/or its regulatory proteins such as Cdc42-GEF^Scd1^, Scd2 or GAP-Cdc42^Rga4^, all of which function at the shmoo site during the mating (Bendezu and Martin, 2013; Dudin et al., 2016)(Fig.4 A and B). In agreement with this hypothesis, a transient MAPK^Spk1^ and MAPKK^Byr1^ signal on the cell cortex was observed during the mating process (Fig. 1)(Dudin et al., 2016). Localisation of MAPK at the growing cell tips was also observed in other fungi including *S. cerevisiae* and *N. crassa* (Chen et al., 2010; Fleissner et al., 2009; Maeder et al., 2007; van Drogen et al., 2001). In budding yeast, MAPK^Fus3^ can directly phosphorylates Bni1, a formin that organises actin filaments, to facilitate shmoo formation (Matheos et al., 2004). It will be important to determine whether this is also the case in fission yeast.

Interestingly, during the vegetative growth when expression of MAPK^Spk1^ is repressed, Ras1.G17V is still capable of activating Cdc42 (Fig. 4C and D). Whether other MAPKs, such as Sty1 or Pmk1, contribute to Cdc42 activation during the vegetative cell cycle will be an important question to answer. Intriguingly, recent studies show that Sty1 inhibits, rather than assists, establishment of the Cdc42 polarity module (Mutavchiev et al., 2016). Collectively, it is likely that the Cdc42 polarity module is regulated in a context dependent manner by multiple MAPKs in a range of ways.

In this study we revealed the vital contribution of Cdc42 to induce the *ras1.G17V* phenotype in fission yeast pheromone signalling (Fig. 8B). In mouse models, small GTPases, Cdc42 and Rac, are required for *H-ras^G12V^* induced transformation (Malliri et al., 2002; Stengel and Zheng, 2012). Therefore, oncogenic RAS-induced Cdc42/Rac misregulation may be a common basis of oncogenicity of mutated-RAS-induced signalling. Specifically targeting this process may therefore be an effective strategy against oncogenic RAS-driven tumourigenesis.

## Materials and methods

### Yeast strains and media

Genotypes of the *Schizosaccharomyces pombe* strains used are listed in the supplementary Table S1. Gene disruption and 2xFLAG-GFP-tagging of genes were performed using the direct chromosomal integration method described previously (Bahler et al., 1998; Funaya et al., 2012; Wach, 1996). Fission yeast media (YE, MM-N and SPA) and basic genetic manipulations are described in (Moreno et al., 1991).

### Plate mating assay for synchronous mating

In order to induce synchronous mating, we established the “Plate mating assay” system as follows: Cells were grown in YE supplemented with adenine (0.2g/L) until the cell density reaches 8-10 x 10^6^ cells/ml. Cells were then washed with MM-N (1% glucose, supplemented with Leucine 40mg/L) using a filtration unit. These cells were resuspended to a cell density of 0.8-1x10^7^cells/ml in MM-N. In the timecourse experiments, this moment was considered time 0. 8x10^7^cells were re-filtered onto a PDVF membrane of 47 mm diameter (Millipore DVPP04700) in order to evenly place the cells across the defined area. The filters bearing the cells over their surface were carefully transferred onto sporulation agar plates (SPA+Leucine 40mg /L, 50ml SPA agar / 14cm diameter for 6 membranes), cell side up, and incubated at 30°C. During the time course experiments, at each time-point one membrane, which initially had 8x10^7^cells at time 0, was removed from the SPA plate and placed into 5 ml of ice-cold 20% trichloroacetic acid (TCA) in a 50 ml falcon tube. The tube was shaken vigorously to remove cells from the membrane. The tube was briefly centrifuged and the membrane removed from the tube. The tube was then centrifuged for 1 minute at 2000rpm to pellet cells. Supernatant was discarded, cells were resuspended in 1 ml 20% TCA and transferred to a tube capable of storage at −80°C. The cells were pelleted again using a bench top centrifuge, supernatant discarded, and the pellet snap frozen in liquid nitrogen. Tubes were stored at −80°C ready for protein extraction.

### Preparation of whole cell extracts

This method is based on that described by Keogh lab Protocols (https://sites.google.com/site/mckeogh2/protocols). All steps were performed on ice unless otherwise stated. Cell pellets were thawed on ice and resuspended in 250 μl 20% TCA. 250 μl of acid washed glass beads (SIGMA) were added and sampled chilled on ice for 5 minutes. Cells were broken in a FastPrep24 cell beater (MP Biomedicals, speed 6.5 x 1 minute x 3 with 1 minute intervals). Cell extracts were collected by piercing the bottom of tubes with a needle and placing them into collection tubes and centrifuged at 2000 rpm for 1 minute. Beads were washed with 300 μl of 5% TCA and centrifuged at 2000 rpm for 1 minute, this was repeated twice. The contents of the collection tube were transferred to a 1.5 ml tube and centrifuged at 14K rpm for 10 minutes at 4°C. The liquid was discarded and the cell pellet washed with 750 μl of 100% ethanol. Tubes were centrifuged briefly (~1min 14000rpm) again. Ethanol was completely removed with a pipette and the pellet resuspended in 100 μl 1M Tris pH8.0 and 100 μl 3x protein loading buffer (Laemmli, 1970). Cell extracts were then heated for 5 minutes at 95°C, centrifuged for 5 minutes at 14K rpm and the supernatant transferred to a fresh 1.5 ml tube. Samples were stores at −20°C.

### Western Blotting

Protein extracts were subject to SDS-PAGE and were transferred to Immobilon-FL PVDF membrane (Millipore). Membranes were blocked for a minimum of 1-hour in Odyssey Blocking Buffer (OBB) (Li-cor) diluted 1:1 in PBS. Primary antibody incubation was carried out overnight in OBB 1:1 in PBS with the anti phospho-p44/42 MAPK (Erk1/2) (Thr202/Tyr204) rabbit monoclonal antibody (#4370 Cell Signaling technology. Used at 1:2000 dilution), the monoclonal anti GFP antibody ((0.4mg/ml), Roche Cat No 11814460001. Used at a 1:2000 dilution) and the anti α-tubulin antibody, TAT1 (generous gift from Keith Gull, 1:3000). Membranes were washed 3x10 minutes with 20 ml TBST (50 mM Tris, pH 7.4 – 150 mM NaCl – 0.1% Tween 20) whilst gently shaking followed by secondary antibody incubation (IRDye 680LT goat anti-mouse antibody, Li-cor 926-32211(1.0 mg/ml), 1:16,000 dilution and IRDye 800CW goat anti-rabbit secondary antibody Li-cor 926-68020 (1.0 mg/ml), 1:16000 dilution in 0.01% SDS, 0.1% Tween 20 in OBB 1:1 PBS) for 1 hour in the dark to prevent fluorophore bleaching. This was followed by 2x 10 minute TBST washes and a 1x 10 minute TBS (50 mM Tris, pH 7.4 – 150 mM NaCl) wash before scanning on the Odyssey CLx infrared Imaging System (Li-cor). Protein quantitation using the Image Studio V2.1 software supplied with the Odyssey CLx scanner. The background method used was median with a border of 3 pixels to the left and right of quantitation boxes.

### Quantitation of MAPK^Spk1^ phosphorylation

To detect the activated MAPK^Spk1^, whole cell extracts were prepared and Western blotting analysis was carried out using the anti-phospho ERK antibody (#4370 Cell Signalling Technology) that recognises the dual phosphorylation at the conserved TEY motif. Specificity of the antibody against the phosphorylated MAPK^Spk1^ (**pp**MAPK^Spk1^) in the fission yeast cell extracts was confirmed before proceeding with further experiments as shown in Supplementary Fig. S1. As described above in the “**Plate mating assay for synchronous mating**”, during the time course experiments, at each time-point, one membrane, which initially had 8x10^7^cells at time 0, was used to prepare the whole cell extracts. Then an equal volume of each sample was loaded to the Western blotting. In this manner, we observed that the change of the protein level throughout the mating process per a defined starting material. We noticed that the level of α-tubulin during the wildtype mating process is relatively constant; less than 25% fluctuation throughout the timecourse for all three biological replicates (an example shown in Fig. S2B). Based on these observations, we chose to use α-tubulin as an internal control to quantitate the **pp**MAPK^Spk1^ signal.

In order to quantitate **pp**MAPK^Spk1^ levels in wildtype, *byr1.DD, ras1.G17V* and *scd1Δ* mutants, Western blotting of three biological replicates of time-course experiments was conducted (original membranes presented in Fig. S3). Because the SDS-PAGE gels we used had only 25 wells, only two out of three biological replicates could be run simultaneously on the same gel. The third replicate was run separately on a different gel as shown in Fig. S3 (A). Definite measurement values of the third replicate were multiplied by a constant to bring the values closest to the rest of the two replicates. In order to directly compare results of wildtype, *byr1.DD, ras1.G17V* and *scd1Δ* mutants, technical replicates of Western blotting were generated by directly comparing one of the biological replicates of each strain as follows: the third replicate of the wildtype and *byr1.DD* samples (WT-3 and *byr1.DD-3*) (Fig. S3A), the second replicate of the wildtype and *ras1.G17V* samples (WT-2 and *ras1.G17V* −2) and the second replicate of the wildtype and *scd1Δ* samples (WT-2 and *scd1Δ* −2) (Fig. 3B).

### Cell Imaging for bright field and Spk1-GFP signals

A Nikon Eclipse Ti-E microscope equipped with CoolLED PrecisExcite High Power LED Fluorescent Excitation System, an Andor iXon EM-DU897 camera and aCFI Plan Apo VC 100x/1.4 objective was used for cell imaging to capture the morphology of cells and the localisation of the GFP-tagged MAPK^Spk1^ during mating time-courses. For cell images presented in Figure 4, a 2D array scanning laser confocal microscope (Infinity 3, VisiTech) was used (see below). Each time point comprises 15-25 serial images with 0.4 μm intervals along the Z axis taken to span the full thickness of the cell. All of the Spk1-GFP images were deconvolved using the Huygens Essential Deconvolution software (Scientific Volume Imaging). Deconvolved images were Z-projected (maximum intensity), cropped and combined using Fiji (Schindelin et al., 2012).

### Imaging and quantitation of CRIB-GFP signal

A 2D array scanning laser confocal microscope (Infinity 3, VisiTech) on a NikonTi-E microscope stand equipped with a Hamamatsu Flash 4.0V2 sCMOS camera and a Plan Apo 100x/1.45 objective was used to capture CRIB-GFP signal to examine activation status of Cdc42. For images presented in Fig. 4A, 25 serial Z images with step size of 0.25 μm were taken. Images were deconvolved using Huygens Essential (Scientific Volume Imaging) and maximum intensity projections are presented.

For images presented in Fig. 4C, for each image, 30-45 serial images with 0.2 μm interval along the Z axis were taken to span the full thickness of the cell. Using FIJI software maximum intensity Z-projections were created and used for quantitation. Intensity of GFP signal on the cell cortex was measured using FIJI along one of the cell tips that shows stronger GFP signal as indicated in an example image in Fig. 4D. 40 cells without septum were measured for each strain. Quantified CRIB-GFP intensity traces were analysed in R (R Core Team http://www.R-project.org/). First, rolling averages over +/−5 measurement points were calculated. Next, the X-coordinates of these traces were aligned by their peak intensity. Finally, the average curve from all aligned traces per strain was calculated, and displayed in Fig 4.D with respective standard error of the mean curves (dashed lines). Analysis script will be accessible on the GiHub repository upon publication (see availability section 1.1.16).

### Glutathione-S-transferase (GST) tag pull-down assays

Fission yeast cDNA fragments encoding Ras1.G17V (1-172), Byr2 (65-180) and Scd1 (760-872) were cloned in the pLEICS1 (Ras1.G17V) and pLEICS4 (Byr2 and Scd1) vectors (Protex [Protein Expression Laboratory], University of Leicester) to produce N-terminally His6-tagged Ras1.G17V (1-172), N-terminally GST-tagged Byr2 (65-180) and N-terminally GST-tagged Scd1 (760-872). Constructs were expressed in *E.coli* BL21 Rosetta cells in LB. All bacteria cell lysates were prepared in phosphate-buffered saline (PBS), pH 7.4.

His6-Ras1.G17V (1-172) was purified using Ni Sepharose 6 Fast Flow (GE Healthcare) and dialysed into buffer A (20 mM Tris-Cl (pH 7.5), 100 mM NaCl, 5mM MgCl2, 1mM β-mercaptoethanol). For GTP loading, buffer A of the Ras1.G17V (1-172) preparation was firstly replaced by Exchange buffer (20 mM Tris-Cl (pH 7.5), 1mM EDTA, 1mM β-mercaptoethanol) using a filtration unit (Amicon Ultra Centrifugal Filter Unit, 10K cut off, Millipore). GDP-GTP exchange reaction was carried out by adding GTP and EDTA to a final concentration of 8 mM and 12 mM respectively and incubating the sample at 37°C for 10 min. The exchange reaction was terminated by adding MgCl2 to a final concentration of 20.5 mM. The resultant GTP-loaded Ras1.G17V (1-172) was washed with buffer A using the filtration unit and immediately used for the GST pull-down assays.

GST-Byr2 (65-180) and GST-Scd1 (760-872) were purified using glutathione sepharose 4B (GE Healthcare). The proteins bound on the glutathione beads were washed with buffer A and used for the GST pull-down assays together with the Ras1.G17V (1-172) preparation. For the competition assays shown in Fig. 7D, Scd1 (760-872) fragment was prepared by cleaving it out from the N-terminal GST, which was bound to the glutathione beads, using TEV protease.

GST pull-down assays were carried out by incubating Ras1.G17V (1-172) (GDP bound or GTP bound) and GST-Byr2 (65-180) or GST-Scd1 (760-872) with or without Scd1 (760-872) fragment at 4 °C for 30 min. Supernatant of the reaction was saved and mixed with the same volume of the 3x loading buffer (Laemmli, 1970) to generate the “unbound” fraction. The remaining beads were washed twice, resuspended in buffer A of the original volume and mixed with the same volume of the 3x loading buffer to generate the “bound” fraction. Protein samples were run on a SDS-PAGE gel and proteins were visualised by InstantBlue protein stain (Expedeon). Quantitation of the intensities of Ras1.G17V (1-172) bands in Fig. 7D was carried out using Fiji software.

### Mathematical Modelling

We built a mathematical model in form of a system of ordinary differential equations (ODE) for the dynamics of the signaling pathway downstream of the pheromone receptor. The model consists of 6 ordinary differential equations representing 9 chemical reactions connecting 6 variables (1.1.14).

We focused on the competition between the MAPK^Spk1^ and Cdc42 pathways for active Ras1^GTP^ molecules, but no other pathway “cross-talks” were instigated. We estimated the model parameters from the measured time courses of **pp**MAPK^Spk1^ in wild type and the three key mutant strains (*ras1.G17V, MAPKK^byr1.DD^* and *Cdc42-GEF^scd1^Δ*) examined in this study. Time courses in these strains are precisely reproduced by the model, thereby we have made it plausible that the competition between MAPK^Spk1^ and Cdc42 pathways is sufficient to cause the **pp**MAPK^Spk1^ early activation phenotype observed in the *ras1.G17V* and *Cdc42-GEF^scd1^Δ* mutants. The model also successfully predicted the **pp**MAPK^Spk1^ temporal dynamics in the 21 other mutant strains examined in this study (Fig.7D). We therefore concluded that the model may serve as a general framework for pheromone signalling in *S. pombe* and can be used for further *in silico* experimental design.

#### 1.1.1 Pathway components

Previous studies showed that more than 20 proteins, more than 20 biochemical and at least 10 relevant genetic interactions are involved in the fission yeast pheromone pathway (Mata and Bahler, 2006; Mata et al., 2002; Otsubo and Yamamoto, 2012; Xue-Franzen et al., 2006). Given the complexity of the pathway, and in order to attain the simplest model fulfilling our criteria, several pathway components and steps were simplified in the presented model as detailed below.

#### 1.1.2 Nitrogen withdrawal, and subsequent pathway activation

Nitrogen starvation induces the expression of the meiotic transcriptional regulator Ste11, which then induces expression of the key components of the pheromone pathway (Egel, 2004). Experimentally, this initial process occurs as a step-like nitrogen removal from the media, which is implemented as a change of starvation signal *S* from 0 →1, i.e. activator step-function in the model.

The response to starvation is two fold. First of all, various signaling components under the regulation of Ste11 are induced: the pheromone (M-or P-factor), the pheromone receptors (Map3 or Mam2), the receptor-coupled G-protein (Gpa1), a GDP-GTP exchange factor (GEF) for Ras1 (Ste6) and an adaptor Ste4. For simplicity, the pheromone sensing and upstream signal transduction machinery is modelled by one variable, called here the Pheromone Sensing (PS) unit (Fig. 7A). The PS unit is sometimes also referred to as the “upstream pheromone pathway”. Induction of the PS unit by Ste11 as well as pheromone binding, subsequent Gpa1 activation and consequential activation of Ste4 and Ste6 are all encompassed in the process [L1] in Fig. 7A. As Ste11 is induced by starvation, PS is also created proportionally to starvation sensing (*S*, Eq. 1). Since downregulation of PS occurs in form of a long negative feedback loop, the term *v_downreguiation_*(*PS*) depends on **pp**MAPK^Spk1^ (Eq. 2); and it is explained in a later section.

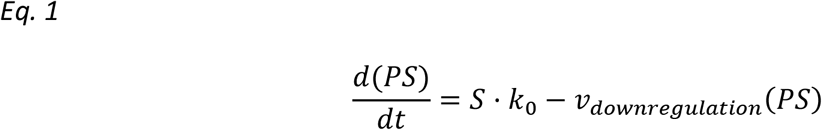

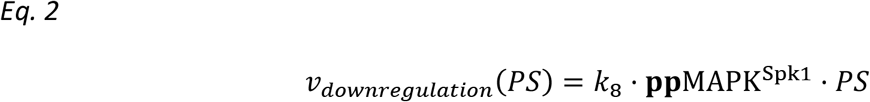

The second response to starvation is the activation of a basal MAPK^Spk1^ transcription through Ste11. MAPK^Spk1^ is then activated by phosphorylation to ppMAPK^Spk1^, if (active) ppMAPKK^Byr1^ is present. This basal activation (F1) is also proportional to starvation sensing (Eq. 3), also see (*Eq. 7*) and section (1.1.6) for details.

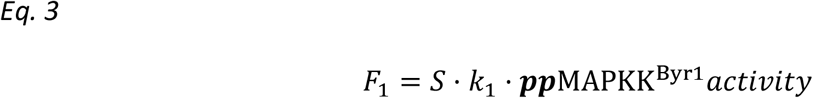

#### 1.1.3 The upstream pheromone pathway (PS)

The PS unit activates the downstream MAPK cascade in two ways:

1. PS activates Ras1^GDP^ (process [L4] in Fig. 7A), which in turn activates MAPKKK^Byr2^, and
2. Ste4, a PS unit component, directly activates MAPKKK^Byr2^ independently of Ras1 (process [L3] in Fig. 7A).

Both of these steps are necessary to functionally activate MAPKKK^Byr2^ (Eq. 6).

Finally, the upstream pheromone pathway is downregulated in a negative feedback loop as discussed in a later section (1.1.7).

#### 1.1.4 Ras1

In the *S. pombe* pheromone response, Ras1 is a central component as it both activates the transcriptional response via MAPKKK^Byr2^, and the morphological response via Cdc42-GEF^Scd1^. Both molecules form protein complexes that anchor pools of Ras1^GTP^ (Chang et al., 1994; Tu et al., 1997; Weston et al., 2013), thus we hypothesised that the two pathway branches might affect each other by changing the amount of available, unbound Ras1^GTP^. In other words, MAPK^Spk1^ and Cdc42 pathways compete with each other for unbound Ras1^GTP^. As the model focuses on the **pp**MAPK^Spk1^ levels, but not on Cdc42^GTP^ production / localisation, the Cdc42 branch (the process [L9]) is simply represented as a change in available Ras1^GTP^ for MAPKKK^Byr2^ in the model. This is further discussed in the section on the *Cdc42-GEF^scd1^Δ* mutant strain (1.1.11).

In the wildtype model, Ras1^GDP^ is activated by the PS unit and deactivated by a Ras-GAP. Both reactions are described by mass action kinetics in (Eq. 4 and Eq. 5). For simplification of parameters, see section (1.1.8).

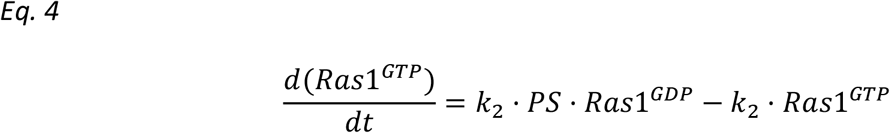

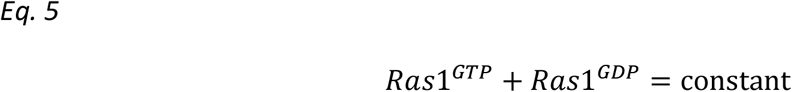

#### 1.1.5 MAPKKK^Byr2^ and MAPKK^Byr1^

Following the PS unit and Ras1^GDP^ activation, MAPKKK^Byr2^ is activated by a twofold mechanism involving Ste4 (a component in the PS unit) and Ras1^GTP^. Note, that MAPK^Spk1^ is the only member of the kinase cascade whose transcription is regulated by Ste11 (the process [L2]) (Mata and Bahler, 2006). Therefore, accordingly, the expression levels of MAPKKK^Byr2^ and MAPKK^Byr1^ were not subject to Ste11 activity in our model. Compared to the 24hrs time scale of pheromone signaling, the MAPKKK^Byr2^ -> MAPKK^Byr1^ activation likely happens on a much faster time scale. Therefore, and for simplicity, we grouped MAPKKK^Byr2^ and MAPKK^Byr1^ into one unit (Fig. 7A). Thus, MAPKK^Byr1^ activation (phosphorylation) (the process [L5]) is given as in (Eq. 6 and *Eq. 7*) and both activation and deactivation follows mass action kinetics. For simplification of parameters, see section (1.1.8).

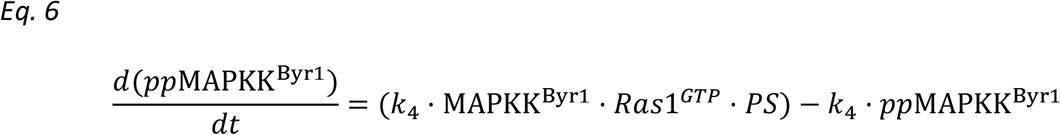

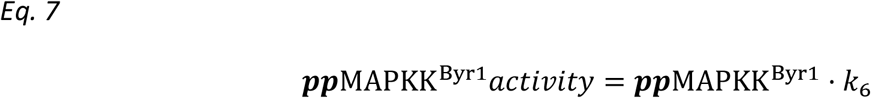

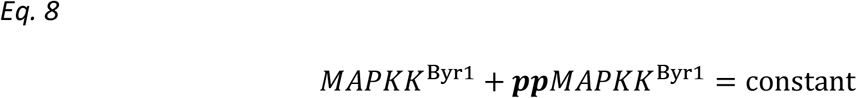

#### 1.1.6 MAPK^Spk1^

In the cells, MAPK^Spk1^ protein expression is induced upon nitrogen removal (the process [L2] in Fig. 7A). MAPK^Spk1^ is then phosphorylated and activated to **pp**MAPK^Spk1^ by **pp**MAPKK^Byr1^ (the process [L6]). Experimentally we observed very similar dynamics for MAPK^Spk1^ and **pp**MAPK^Spk1^. Therefore, we assumed that MAPK^Spk1^ ~ **pp**MAPK^Spk1^ system is in equilibrium, so creation and destruction reactions for each state can be disregarded. Thus, for simplicity, MAPK^Spk1^ is only present in its phosphorylated form (**pp**MAPK^Spk1^) in the model.

Despite the simplification, the model represents the three main processes contributing to **pp**MAPK^Spk1^ levels: it is created proportionally to starvation and to MAPKK^Byr1^ activity (F_1_, *Eq. 3*); it is also self-induced (which is again proportional to MAPKK^Byr1^ activity (F_6_, *Eq. 7-Eq. 9*); and it is degraded by mass action kinetics (*Eq. 11*).

##### 1.1.6.1 Positive Feedback

Previous studies have shown that **pp**MAPK^Spk1^ activates Ste11 (Kjaerulff et al., 2005), which in turn transcribes more *mapk^spk1^*. This forms a positive feedback loop (the process [L7]) if **pp**MAPKK^Byr1^ is present to phosphorylate MAPK^Spk1^. A simplest form of an abstract positive feedback is given as the self-dependence term **pp**MAPK^Spk1^*k*_9_^^ in Eq. 9.

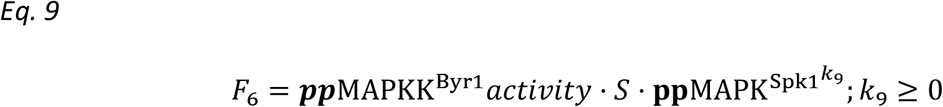

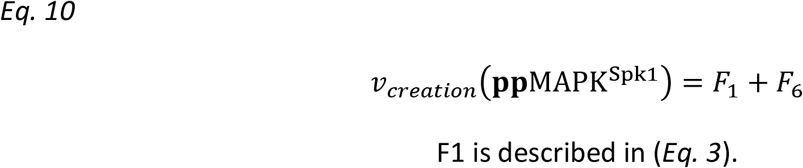

Finally, MAPK^Spk1^ is set to be degraded following simple mass action kinetic, thus the final equation for ppMAPK^Spk1^ kinetics in wild type yeast is given as (Eq. 11).

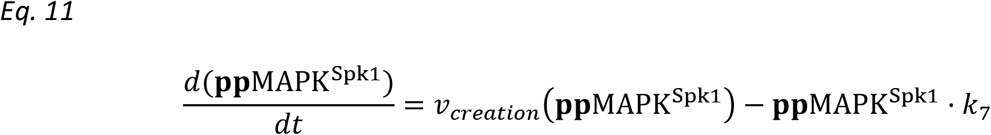

#### 1.1.7 Down regulation of the pathway by a negative feedback loop

Transient MAPK^Spk1^ activation in the wildtype cells (Fig.1C) suggests that there is a delayed negative feedback to downregulate the **pp**MAPK^Spk1^ in the pathway, which leaves only a certain time-window for activity. As this downregulation was found abolished in the constitutively active *MAPKK^byr1.DD^* mutant (Fig. 1E), the target(s) of the negative feedback must be MAPKK^Byr1^ or upstream components. Hence, potential negative regulators, such as phosphatases Pyp1 and Pmp1 that are proposed to directly deactivate ppMAPK^Spk1^ (Didmon et al., 2002), were excluded from our model. Meanwhile, the negative regulators Sxa2 (encoding a serine carboxypeptidase against a mating pheromone P-factor) and Rgs1 (encoding a regulator of Gpa1), both of which are transcriptionally induced upon pheromone signalling, fit well to the criteria of a negative feedback component (Imai and Yamamoto, 1992; Mata and Bahler, 2006; Pereira and Jones, 2001; Watson et al., 1999). Receptor internalization is also expected to be a part of the negative feedback (Hirota et al., 2001) and regulation by antisense RNA for the *mapk^spk1^* transcript as well as other signalling components may also play a role (Bitton et al., 2011). As relative contributions of each remain to be quantified, we represented all these potential negative feedbacks collectively as a single circuit in our model that inactivates the PS unit (the process [L8]) (Fig 7A). The chemical equation is therefore given as: *Ras1^GTP^* + *ppMAPK*^*Spk*1^ + *PS* → *Ras1^GTP^* + *ppMAPK^Spk1^*, which gives rise to the negative term in Eq. 12.

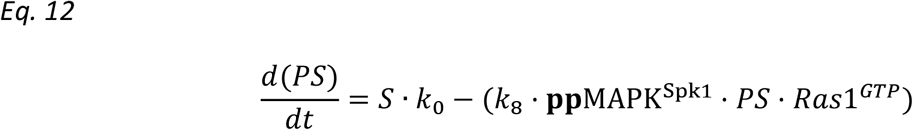

##### 1.1.7.1 Ras1^GTP^ contribution to the negative feedback

Transient, peaking ppMAPK^Spk1^ activation in wildtype solely requires a delayed negative feedback. However, as described in the main text, a more complex regulation is necessary to describe the both earlier and lower peak in *ras1.G17V* and *cdc42-GEF^Scd1^Δ* mutants. By incorporating Ras1^GTP^ in this feedback (the process [L10], see: Eq. 12), we could precisely recapitulate both phenotypes.

##### 1.1.7.2 The “simple-activator-Ras” model variant

We wanted to show the behaviour of the model if Ras1^GTP^ would only act as an activator. From the wild type model presented in Fig.7 we removed Ras1^GTP^ from the downregulation resulting in (Eq. 13).

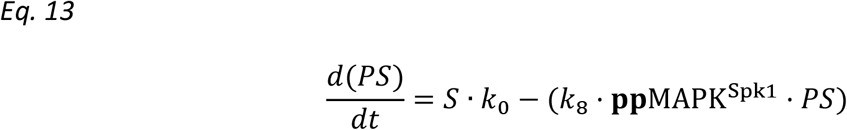

We fitted this model variant to the data sets as before, using Hooke–Jeeves (Hooke and Jeeves, 1961) and particle swarm optimisation methods (Kennedy and Eberhart, 1995). The objective value for the best fit was ~3 higher for this model variant than for the original model. Although it recapitulated the main features (Fig.S7B), it could not describe the observed “earlier and lower” signalling phenotype observed in the *ras1.G17V* and *cdc42-GEF^Scd1^Δ* strains. Finally, we also show that the addition of Ras1^GTP^ to this model only leads to increased amplitude, not to earlier timing (Fig.S7C).

#### 1.1.8 Model simplification

After extensive parameter estimations, we found that the model can be simplified in the number of individual parameters for forward and reverse reaction of the same species without compromising the quality of fit. For the Ras1^GDP^ ←→ Ras1^GTP^ and MAPKK^Byr1^ ←→ ppMAPKK^Byr1^ reactions we could set the forward and reverse parameters to the same value, and achieve virtually identical quality fits, while reducing the number of free dimensions in the parameter space by two. With this simplification the remaining model parameters became uniquely identifiable, with the exception of k1 (Fig.S6). We also fitted the models with these as free parameters and achieved very similar fit qualities, but most parameters became unidentifiable, therefore, we decided for the simpler model.

### Mutant strains

#### 1.1.9 MAPKK^byr1.DD^

MAPKK^Byr1.DD^ harbors two phosphomimetic point mutations, which keep MAPKK^Byr1.DD^ active even in the absence of its activator MAPKKK^Byr2^ (Fig. 3). Therefore, this mutation effectively uncouples MAPK^Spk1^ from the upstream part of the pathway, as represented in Fig. 7A, *MAPK^byr1.DD^* panel. In this mutant, MAPK^Spk1^ (its substrate) also becomes constitutively phosphorylated and activated (Fig. 1E).

It is known that the phosphomimetic substitution of the amino acids often does not fully “mimic” the phosphorylated status and often fails to bring a full activation of the modified kinase. This is likely happening to the MAPKK^Byr1.DD^ mutant molecule: the initial production of **pp**MAPK^Spk1^ upon nitrogen starvation was much slower in the *MAPKK^byr1.DD^* mutant than in wildtype (Fig. 1E). This effect is represented by the *f_Byr1DD_* <1 fitted parameter in the MAPK^Byr1.DD^ model variant (see: Eq. 14).

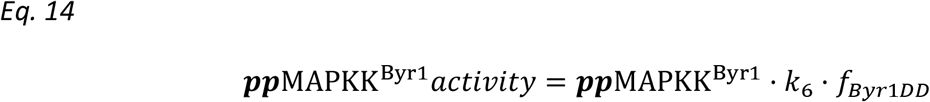

As MAPK^Byr1^ is not induced upon starvation (Nadin-Davis and Nasim, 1990), in the mutant strain, we could implement the mutation by setting initial concentrations [*Byr*1]_0_ = 0 and [*Byr*1^*PP*^]_0_ = 1; and setting the respective activation and inactivation reactions to zero.

#### 1.1.10 ras1.G17V

The *ras1.G17V* strain contains a point mutation, equivalent to human Ras G12V oncogenic mutation, which impairs the GTPase activity of the molecule (Trahey and McCormick, 1987). This results in being most of the molecules to stay in the Ras1^GTP^ form.

In the model, the following changes are implemented to represent the mutation: The initial concentration of [*Ras*1^*GTP*^]_0_ is set to 1, while that of [*Ras*1^*GDP*^]_0_ is set to 0, which is the opposite of the wildtype initial state. Also, the deactivation and activation reactions are set to zero.

#### 1.1.11 Cdc42-GEF^scd1^Δ

Ras1^GTP^ is proposed to interact with and activate Cdc42-GEF^Scd1^, which then activates Cdc42 by acting as a GDP-GTP exchange factor (GEF) (Chang et al., 1994). Because Cdc42-GEF^Scd1^ binds Ras1, the loss of it leads to an increased pool of unbound Ras1^GTP^. In our model this mutation is implemented as additional, unbound Ras1^GTP^ added to the wildtype system. In this setting the amount of Ras1^GTP^ available for MAPKKK^Byr2^ determines the **pp**MAPK^Spk1^ dynamics. Since it is unknown, how much Ras1^GTP^ is bound by Cdc42-GEF^Scd1^, we fit the [*Ras*1^*GTP*^]_0_ as a free parameter in the otherwise intact, wild type model.

#### 1.1.12 Implementation and availability

The **pp**MAPK^Spk1^ temporal dynamics in the four strains, wildtype, *MAPKK^byr1.DD^, ras1.G17V* and *cdc42-GEF^Scd1^Δ*, were described by 6 ODEs and 8 parameters shared across all species (List of Differential Equations, Supplementary Table S2). *MAPK^byr1.DD^* has an additional parameter, *f_Byr1DD_*, while [*Ras*1^*GTP*^]_0_ is fitted the *Cdc42-GEF^scd1^Δ* strain as described above. The model represents relative concentrations of proteins on an absolute time scale. Initial concentrations of components are therefore either set to 1, if the component is present prior to starvation, or 0, if the component is not present, and it is induced or activated during starvation or pheromone response. To show that the model is at steady state before nitrogen removal, it was simulated for 100 hours prior to starvation in every case. All variables remained constant during this time. All other plots start from T=t-100 as origin, when nitrogen is removed.

Model simulation and parameter estimation (model fitting) was performed in COPASI 4.20 (Hoops et al., 2006). Visualisation and data analysis was done in python v2.7 and R v3.2. Plotting was performed by base R graphics and by the MarkdownReports v2.9.5 (Vertesy, 2017), pheatmap (Kolde, 2013) and plotly v4.7.1 (Sievert et al., 2017) packages.

#### 1.1.13 Parameter Estimation

Parameter estimation consisted in fitting of 8 rate constants (parameters) and one additional parameter per mutant strain. We used the gradient-free Hooke–Jeeves (Hooke and Jeeves, 1961) and particle swarm optimisation methods (Kennedy and Eberhart, 1995) implemented in COPASI.

We found that the algorithm converges better to the optimal solution, if key time points in the dataset are counted with different weights for optimisation. Therefore, data points 102 (early activation), 108 and 109 (peak) were passed on with triple weight to the optimisation function.

##### 1.1.13.1 Parameter identifiability, multiplex parameter estimation and ensemble modelling

To investigate parameter identifiability, we performed multiplexed parameter estimation by 5000 parallelised particle swarm optimisation. In each of the estimation rounds, the model was fitted to the data, starting from randomised parameters. Parameters were sampled uniformly +/− 4 orders of magnitudes around the previously achieved best fit, except the exponent (N). This is the exponent of the positive feedback loop, is therefore an ‘explosive’ parameter, and there is no reason to assume a stronger dependence than 4-th order (x^4^) for **pp**MAPK^Spk1^ in the positive feedback. The range for this parameter was therefore set to [1e-5, 4].

We repeated the estimations with +/− 5 magnitudes (5000 times), but we achieved no better fit, while the convergence of optimisation dramatically decreased (i.e. >90% of the optimisations did not get close to the optimal fit).

We found that with increasing fit quality (lower objective value) all parameters except k1 converge to a single value, or a very limited range (Fig.S6D, top row, objective value <5).

In Fig.S7A, we further show the convergence on linear scale for an extended number of simulations (objective value <8). The convergence of parameters is even more pronounced. In suboptimal fits (O.V. >>3.6) [*Ras*1^*GTP*^]_0_ in the *scd1Δ* strain frequently hits the upper boundary, which marks a 10-fold increase in available Ras1^GTP^. Although the boundary is arbitrary, it is unreasonable to think that there will be a bigger increase to the available Ras1^GTP^ pool by the knockout of *scd1*.

Multiplexed parameter estimation for the simple-Ras model performed in the same manner as for the wildtype, 1000 fits with particle swarm optimisation from randomly sampled starting points from a parameter space +/− 4 magnitudes around the best previous fit.

##### 1.1.13.2 Ensemble modelling

To show the reliability of the model dynamics in spite of the un-identifiability of certain parameters, we applied ensemble modelling (Brannmark et al., 2010). In this approach, we analysed the predicted time courses of the 100 best fitting models to show that regardless of the residual parameter uncertainty, the simulated variables’ dynamics are robustly predicted (Fig. S8). To show the variation of the individual simulation, ten thousand randomly sampled simulation points are displayed behind the running average.

#### 1.1.14 List of Differential Equations

As introduced above, here listed and separately defined for all species. Equations 4b and 6b are only listed here, and provided for completeness: they are simply the reverse reactions for 4 and 6 respectively, thus not discussed in the section above.

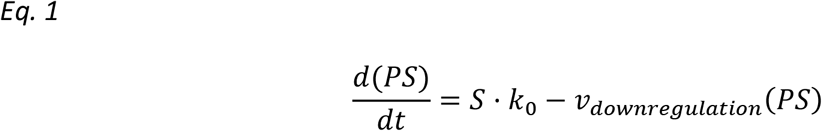

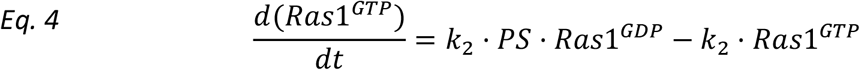

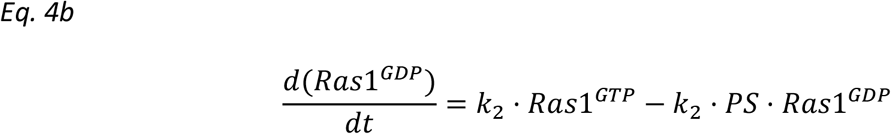

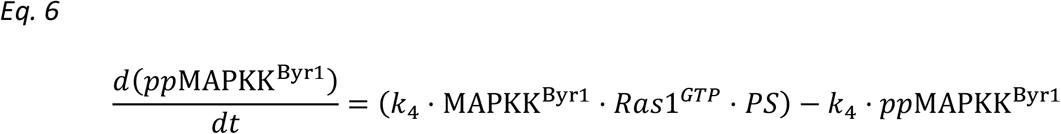

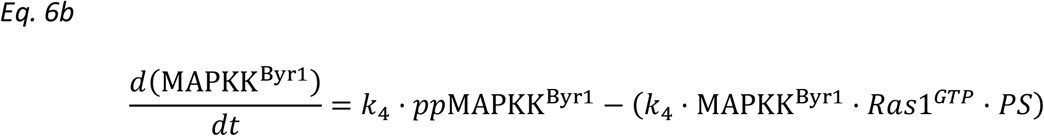

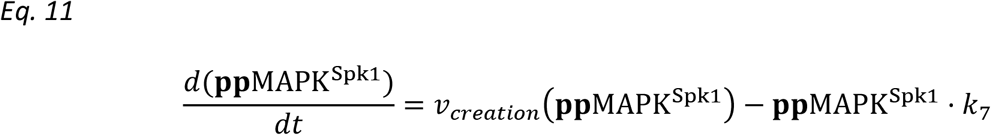

#### 1.1.15 List of Algebraic Equations

(All introduced above)

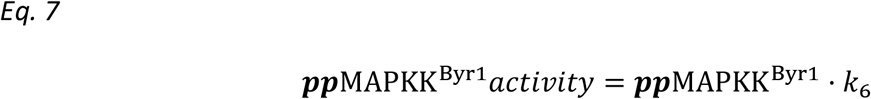

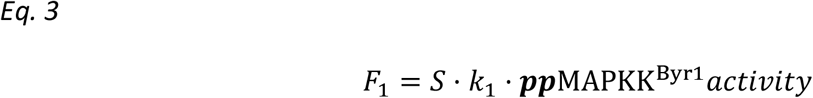

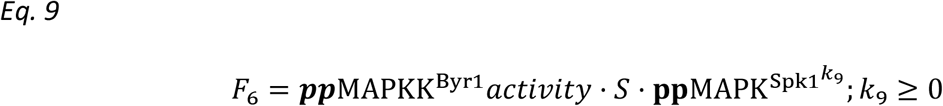

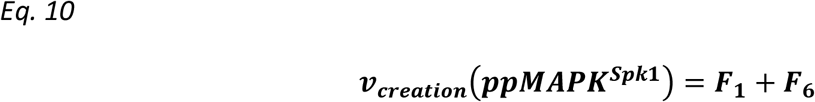

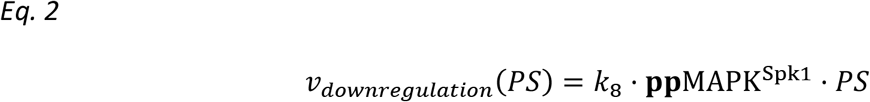

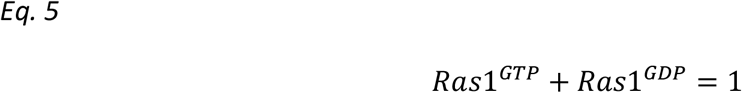

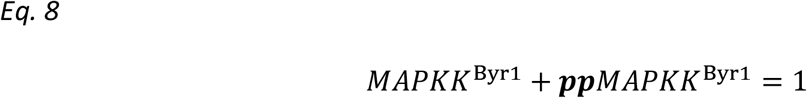

#### 1.1.16 Availability

Upon acceptance, the source code for analysis is available “as-is” under GNU GPLv3 at https://github.com/vertesy/Competition.Model.of.Ras.in.Pombe.Pheromone.Response. The model file will be provided at https://www.ebi.ac.uk/biomodels-main/ in SBML “Level 3: Version 1” format exported from COPASI. The recommended COPASI format model file will be available at the GitHub repository.

## Acknowledgment

The authors thank Tatsuya Maeda, Thibault Mayor, Janni Petersen, Louise Fairall, John Schwabe, David Critchley, Andrey Reviyakin, Gary Willars and Mohan Harihar for helpful suggestions, stimulating discussions and critical reading of the manuscript. We thank PROTEX and PNACL at University of Leicester for their technical assistance. We are grateful to Kazu Shiozaki, Keith Gull and Yeast Genetic Resource Center for providing strains and antibodies. This work was funded by the Wellcome Trust Institutional Strategic Support Fund WT097828/Z/11/Z, WT097828/Z/11/B and the Deutsche Forschungsgemeinschaft (DFG) EXC 81. A.V. was supported by German Academic Exchange Service (DAAD) A0981674.

## Author Contributions

E.J.K., S.R. and K.T. generated yeast strains. E.J.K., G.S. and K.T. monitored activation status of MAPK^Spk1^ and Cdc42. A.V. and E.K. conducted mathematical modelling. E.J.K., K.S. and K.T. conducted image analysis. M.T., R.G., C.P. and C.D. conducted biochemical analysis of Ras1, Byr2 and Scd1. E.J.K., A.V., E.K. and K.T. designed the experiments and interpreted the data.

## Supplementary Information

### Supplementary Figures

**Supplementary Figure S1.**
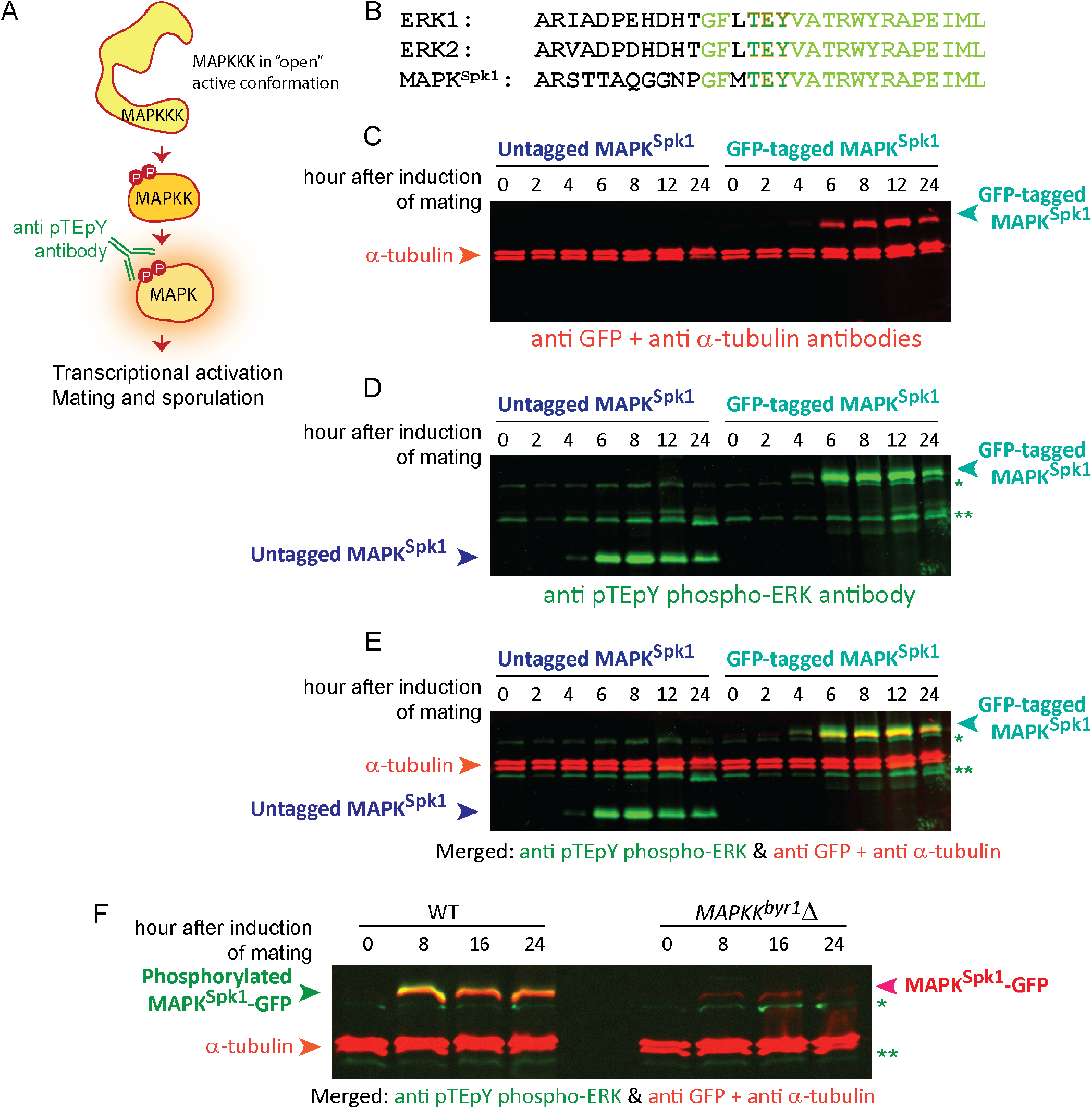
Validation of the use of a commercially available anti-GFP and anti-phospho-MAPK specific antibody. (A) Schematic outlining of the principal of a phospho-specific MAPK antibody, which specifically recognize the phosphorylated TEY motif (pTEpY) of MAPK. (B) Sequence alignment between mammalian ERK1, ERK2 and *S.pombe* MAPK^Spk1^ at the region of the dual phosphorylation required for activation and surrounding residues. Conserved residues are in light green and TEY is highlighted with orange-green. (C)-(E) A Western blot of a time-course of a WT (KT301) and a MAPK^Spk1^-GFP strain (KT3082) over 24 hours, incubated with anti GFP, anti-pTEpY phospho-ERK and anti α-tubulin antibodies. Li-cor Odyssey system was used to detect the signals. (C) The signals obtained by the 700 nm wavelength scan to detect signals from a primary monoclonal mouse anti-GFP antibody ((0.4mg/ml), Roche Cat No 11814460001. Used at a 1:2000 dilution) and the primary anti α-tubulin antibody TAT1 (generous gift from Keith Gull, 1:3000), followed by the IRDye 680LT secondary antibody (goat anti-mouse antibody, Li-cor 926-32211(1.0 mg/ml), 1:16,000 dilution). α-tubulin was used as an internal loading control. (D)The exact same blot as in (C) but showing the 800 nm wavelength scan to detect signals from an anti pTEpY phospho-ERK antibody (#4370 Cell Signaling technology. Used at 1:2000 dilution) visualised by the IRDye 800CW goat anti-rabbit secondary antibody (Li-cor 926-68020 (1.0 mg/ml), 1:16000 dilution). The band indicated by a single asterisk (*) was concluded to be irrelevant to MAPK^Spk1^-GFP because although this band runs very close to MAPK^Spk1^-GFP, it does not exactly overlap with the MAPK^Spk1^-GFP signal (in red) in (E) and (F). Furthermore, the band exists in non-tagged MAPK^Spk1^ strain seen in (D) and in *mapkk^byr1^Δ* strains in (F), further supporting that the band is irrelevant to phospho-MAPK^Spk1^-GFP. A double asterisk indicates a band which was also concluded not to be relevant to phospho-MAPK^Spk1^ because of the same reason. Considering the molecular weight, this double asterisk band may correspond to another MAPK, MAPK^Pmk1^. (E) An overlay of the 700 and 800 nm channels presented in (C) and (D). (F) The phospho-ERK antibody is phospho-specific. Cell extracts were prepared from the time-course of MAPK^Spk1^-GFP tagged WT (KT3082) and *mapkk^byr1^Δ* strains (KT4300) and MAPK^Spk1^-GFP was detected by anti-GFP (red) while phospho-MAPK^Spk1^ was detected by anti-pTEpY phosphor ERK antibody (green). In the *mapkk^byr1^Δ* strain, phosphorylated MAPK^Spk1^-GFP signal is missing although MAPK^Spk1^-GFP was detectable at 8, 16 and 24 hours after induction of mating. The same background bands seen in (D) and (E) are indicated in the same way using a single (*) and double (**) asterisk.

**Supplementary Figure S2.**
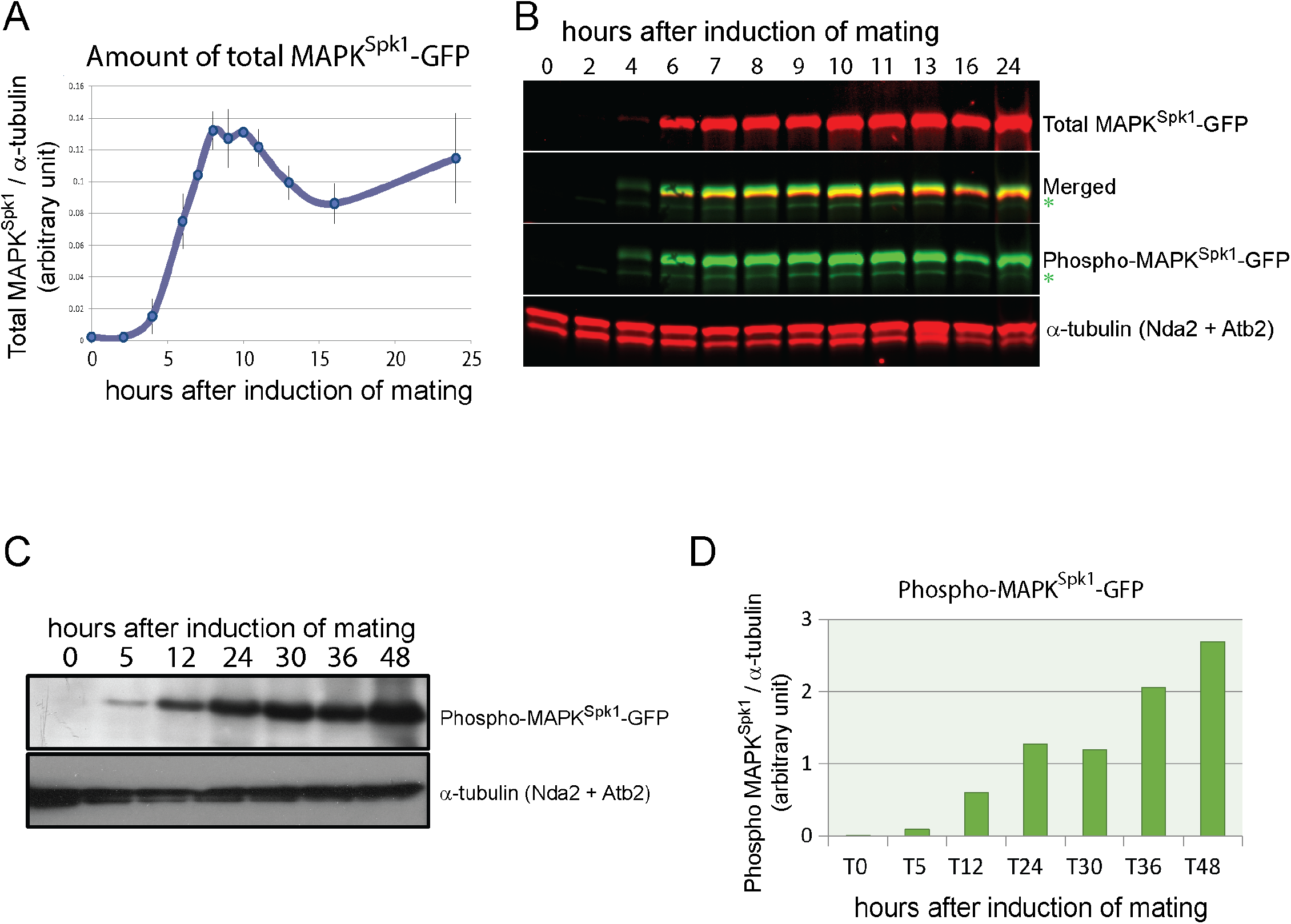
(A) and (B): Quantitation of total MAPK^Spk1^-GFP during the mating process. The total MAPK^Spk1^-GFP protein level was detected using the anti-GFP antibody ((0.4mg/ml), Roche Cat No 11814460001. Used at a 1:2000 dilution). MAPK^Spk1^-GFP Cells (KT3082) were induced for sexual differentiation by the plate mating assay system as described in the materials and methods. (A) Three biological replicates were used for quantitation to obtain the mean value (error bars are ±SEM). α-tubulin was used as a loading control and quantitation was carried out using the Image Studio ver2.1 software (Licor Odyssey CLx Scanner). (B) A typical example of Western blotting used for quantitation is shown. The green asterisk indicates the background band recognized by anti-phospho-ERK antibody (#4370 Cell Signaling technology. Used at 1:2000 dilution) as seen in the supplementary Figure S1. **(C) and (D): *MAPKK^byr1.DD^* mutation causes constitutive activation of MAPK^Spk1^**. Cells carrying *MAPKK^byr1.DD^* mutation (KT3435) were induced for sexual differentiation by the plate mating assay system as described in the Materials and Methods. Western blotting of phospho-MAPK^Spk1^ and α-tubulin was conducted using ECL system (GE healthcare). (A) The results were visualized by exposing the signals to a X-ray film. (B) The bands were scanned and quantitated using ImageJ software (https://imagej.nih.gov/ij/). The α-tubulin signals were used as an internal control to estimate relative ppMAPK^Spk1^ levels.

**Supplementary Figure S3.**
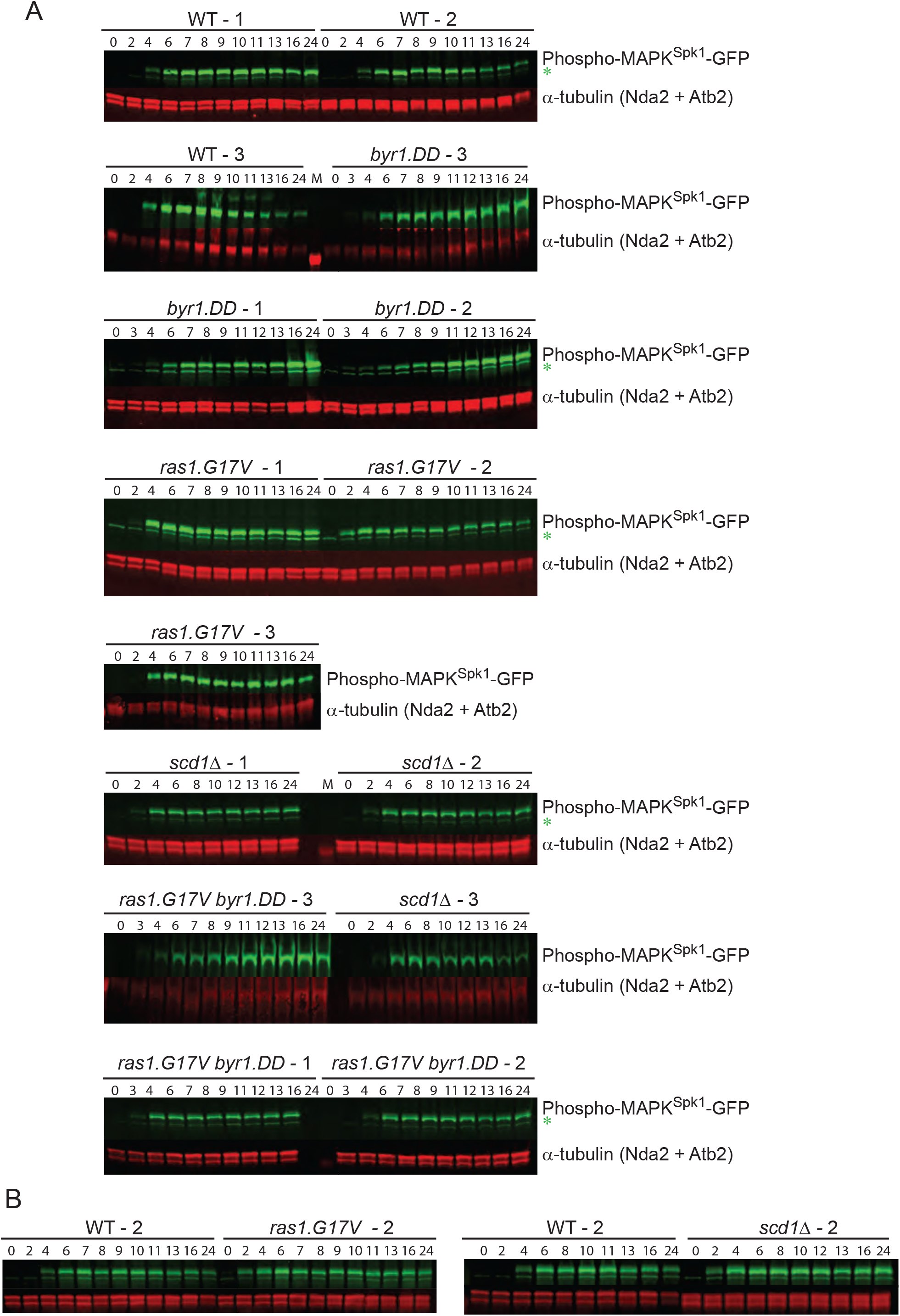
Original Western blotting images of WT, *ras1.G17V, byr1.DD, ras1.GVbyr1.DD* and *scd1Δ* mutants used to quantitate ppMAPK^Spk1^-GFP during the mating process in Fig. 1, 2 and 3. MAPK^Spk1^-GFP Cells were induced for sexual differentiation by the plate mating assay system as described in the materials and methods. (A) Three biological replicates presented were used for quantitation to obtain the mean value and SEM. α-tubulin was used as a loading control and quantitation was carried out using the Image Studio ver2.1 software (Licor Odyssey CLx Scanner). **pp**MAPK^Spk1^-GFP was detected by anti-phospho-ERK antibody (#4370 Cell Signaling technology. Used at 1:2000 dilution). The green asterisk indicates the background band as seen in the supplementary Figure S1. Numbers indicate hours after induction of mating. M: Marker lane. (B) Extracts of WT −2 and *ras1.G17V* −2 (left panel) and WT-2 and *scd1Δ* −2 (right panel) were re-run as technical replicates. These membranes were used to estimate relative **pp**MAPK^Spk1^ intensities between WT, *scd1Δ* and *ras1.G17V* strains.

**Supplementary Figure S4.**
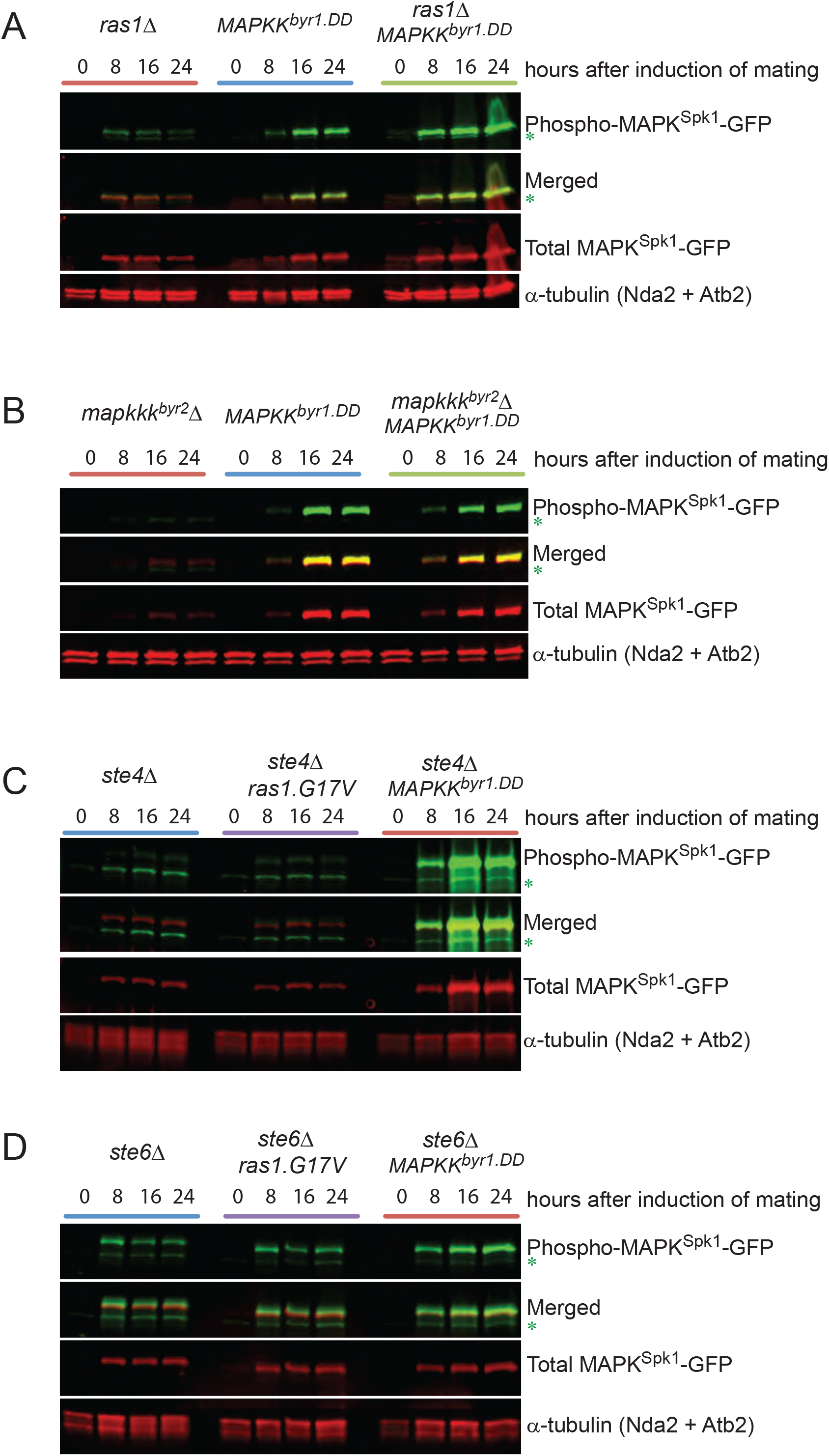
Original Western blotting membranes used in Fig. 3 and 5. (A) Original Western blotting membrane used to quantitate phospho-MAPK^Spk1^-GFP to generate the graph presented in Figure 3 (D). Strains used were; *ras1Δ* (KT4323), *MAPKK^byr1.DD^* (KT3435) and *ras1Δ MAPKK^byr1.DD^* (KT4359) (B) Original Western blotting membrane used to quantitate phospho-MAPK^Spk1^-GFP to generate the graph presented in Figure 3 (E). Strains used were; in *mapkkk^byr2^Δ* (KT3763), *MAPKK^byr1.DD^* (KT3435) and *mapkkk^byr2^Δ MAPKK^byr1.DD^* (KT4010). (C) Original Western blotting membrane used to quantitate phospho-MAPK^Spk1^-GFP to generate the graph in Figure 5 (A). Strains used were; *ste4Δ* (KT4376), *ste4Δ ras1.G17V* (KT5143) and *ste4Δ MAPKK^byr1.DD^* (KT5136). (D) Original Western blotting membrane used to quantitate phospho-MAPK^Spk1^-GFP to generate the graph in Figure 5 (C). Strains used were; *ste6Δ* (KT4333), *ste6Δ ras1.G17V*(KT4998) and *ste6Δ MAPKK^byr1.DD^* (KT5139). Indicated mutant cells were induced for mating by the plate mating assay system as described in the Materials and Methods. Western blotting of phospho-MAPK^Spk1^ and α-tubulin was conducted in the same way as Supplementary Figure S1. The green asterisks indicate the same background band recognized by anti-phospho-ERK antibody (#4370 Cell Signaling technology. Used at 1:2000 dilution) seen in the Supplementary Figure S1 and do not overlap with the red signals that represent the “Total MAPK^Spk1^-GFP” in the “Merged” panel. α-tubulin was used as a loading control and quantitation was carried out using the Image Studio ver2.1 software (Licor Odyssey CLx Scanner).

**Supplementary Figure S5.**
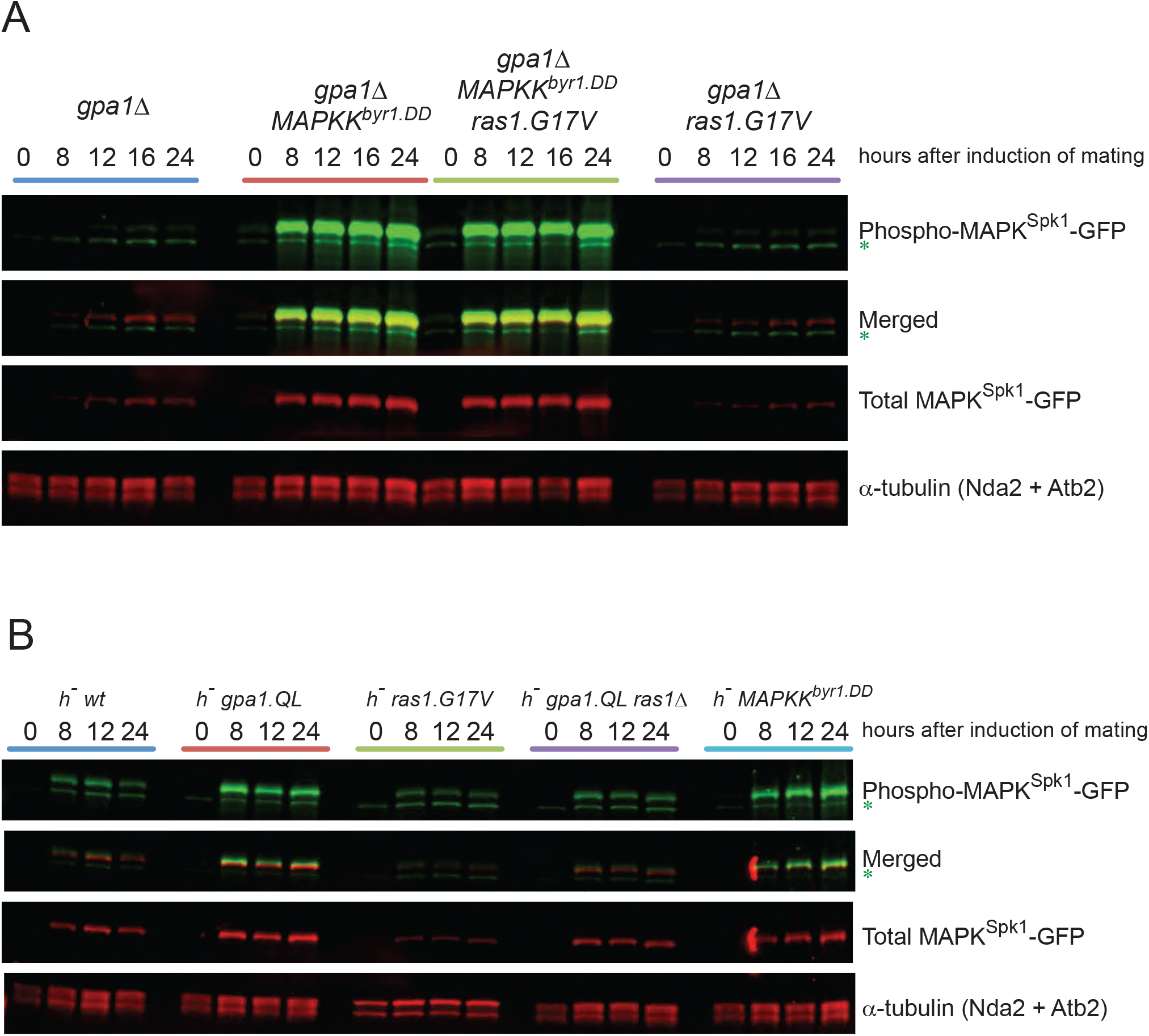
Original Western blotting membranes used in Fig. 6. (A) Original Western blotting membranes used to quantitate phospho-MAPK^Spk1^-GFP to generate the graph in Figure 6 (A) Strains used were; *gpa1Δ* (KT4335), *gpa1Δ ras1.G17V* (KT5023), *gpa1Δ MAPKK^byr1.DD^* (KT4353) and *gpa1Δ ras1.val17 MAPKK^byr1.DD^.(KT5035)*. (B) Original Western blotting membrane used to quantitate phospho-MAPK^Spk1^-GFP to generate the graph in Figure 6. (C) Strains used were; *h^−^* WT (KT4190), *h^−^ gpa1.QL* (KT5059), *h^−^ ras1.G17V* (KT4233), *h^−^ gpa1.QL ras1Δ* (KT5070), h^−^ *MAPKK^byr1.DD^* (KT4194). The green asterisk indicates a background band recognized by anti-phospho-ERK antibody (#4370 Cell Signaling technology. Used at 1:2000 dilution). α-tubulin was used as a loading control and quantitation was carried out using the Image Studio ver2.1 software (Licor Odyssey CLx Scanner).

**Supplementary Figure S6.**
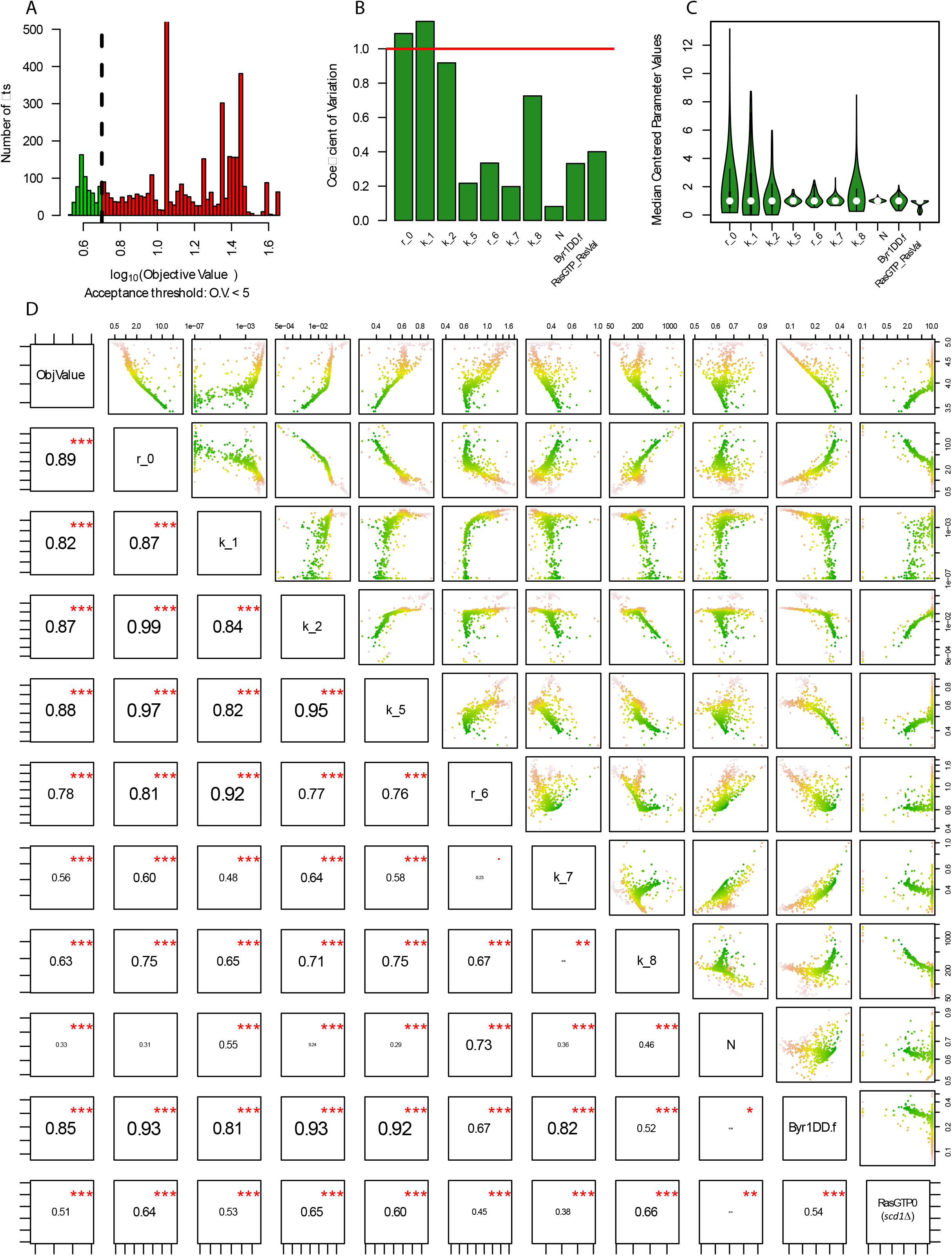
Parameter distributions and relationships in multiplexed global parameter fitting. (A) Distribution of objective values after 5000 parallel parameter estimations by the global optimization algorithm “particle swarm” with randomized starting values within +− 4 magnitudes around the previously achieved best fit. Fits with an objective value < 5 are accepted for further analysis (green). Simulations with +− 5 magnitudes resulted in the same best fits, but much smaller convergence (See Materials and Methods). (B-C) Coefficient of variation and distribution of parameters within the 578 the accepted fits show unimodal distributions for all parameters, and most of them are narrowly distributed, suggesting a unique optimum. In (C), the only bimodal distribution is that of RasGTP0 (Scd1Δ) among fits below O.V.<5. In (D, top right corner) however it is shown that this parameter also converges to a single optima, below O.V. of 3. (D) Pairwise scatterplots and Spearman correlation coefficients among all parameters and the objective value (log scale axes). In this figure, every dot is an independently fitted model (obj. value <5); and every panel shows a pairwise correlation between a fitted parameter and the objective value (top row), or between two fitted parameters (other rows). Green to brown denotes the objective values between 3.4 and 5. The top row shows that with better fits, all parameters (but k_1) converge to a single optimum (bottom most points, in green).

**Supplementary Figure S7.**
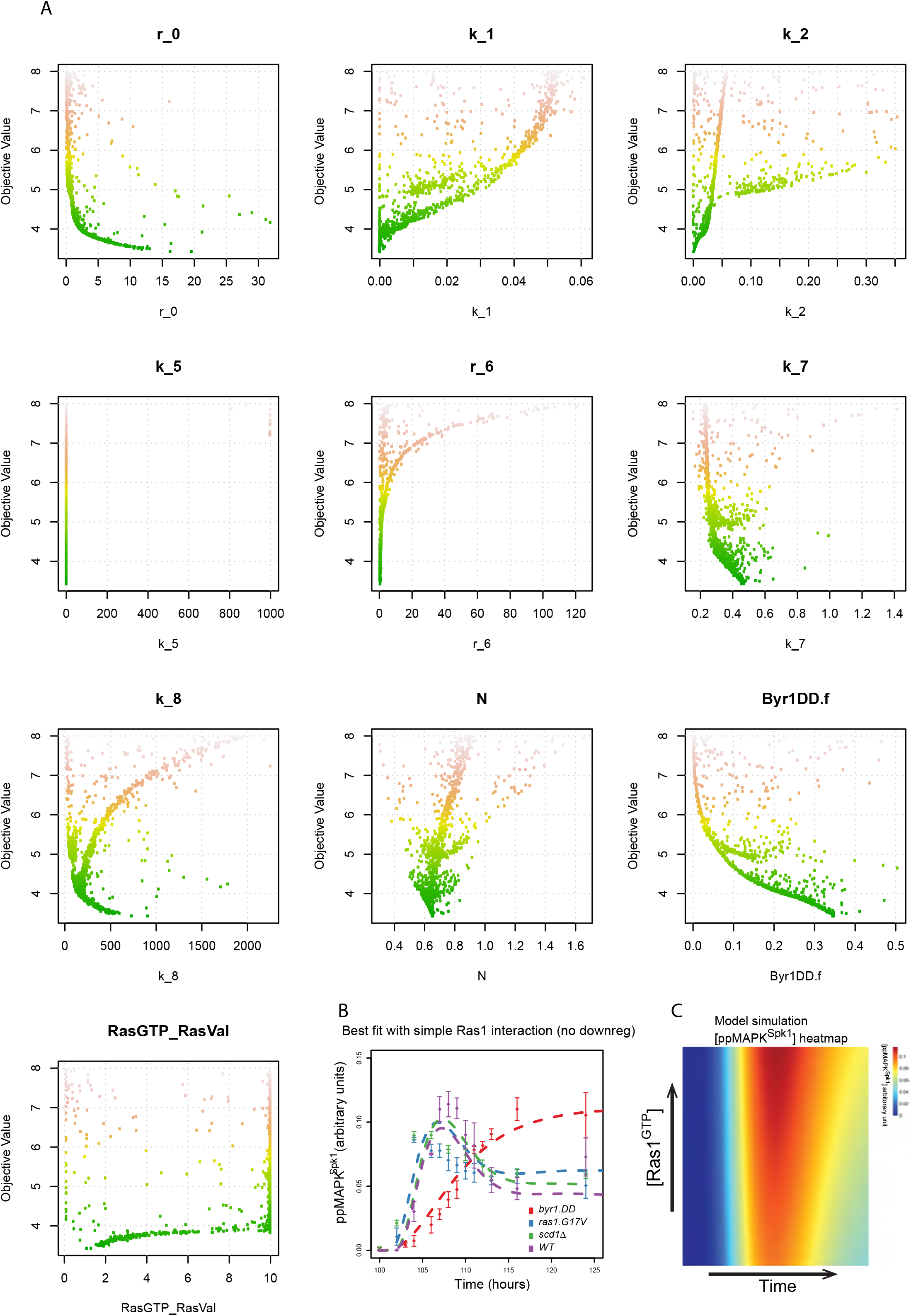
Convergence of parameters in linear scale, over an extended range. (A) Alternative representation of the top row of Fig.S6D. Parameter values are displayed on a linear scale (X-axis) and fits below and objective value (O.V.) of 8 (instead of below 5) are displayed. (B) Simulated time course of MAPK^Spk1^ in the best fit of a simplified model variant shows no advanced peaking in the *ras1.G17V* and *scd1Δ* mutant strains. This model is simplified from the original model (Fig.7 A) by removing [L10], and models the scenario where Ras has no complex interactions, it has only activating roles [L5, L9]. (C) Simulated ppMAPK^Spk1^ dynamics in the simplified model (B) at increasing concentrations of Ras1^GTP^ added *in silico* to the system. Increased Ras1^GTP^ concentration causes advanced and reduced **pp**MAPK^Spk1^ peak intensities.

**Supplementary Figure S8.**
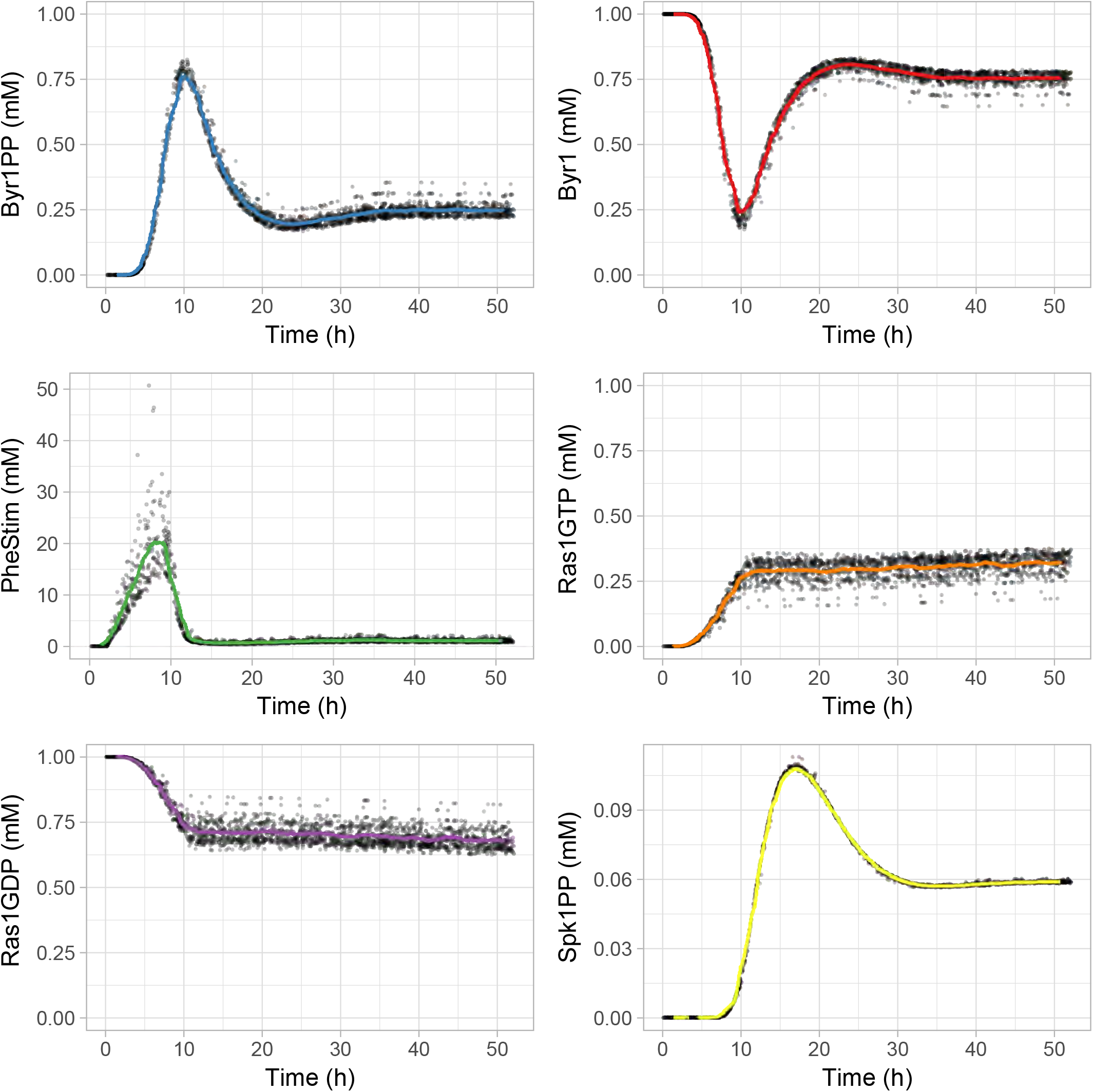
Ensemble modeling shows that the dynamics of protein concentration are robustly predicted regardless of certain parameters’ unidentifiability. Time course simulation of the 100 best-fitted wild type models for all 6 variables. Rolling averages (window of 100) are displayed in color, and ten thousand randomly sampled simulation points (in grey) for each experiment show a narrow distribution of predicted concentration dynamics.

**Supplementary Table S1:**
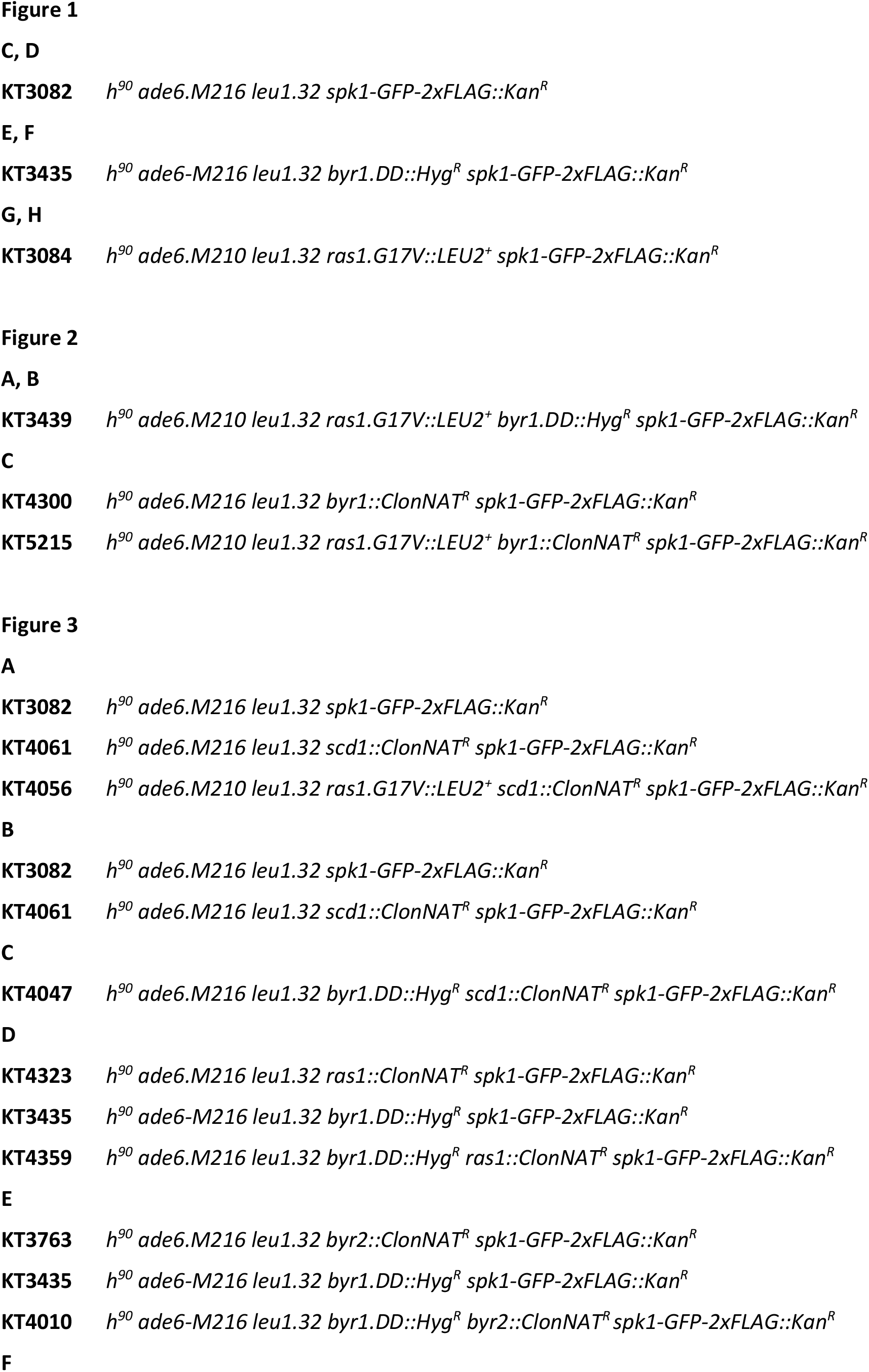
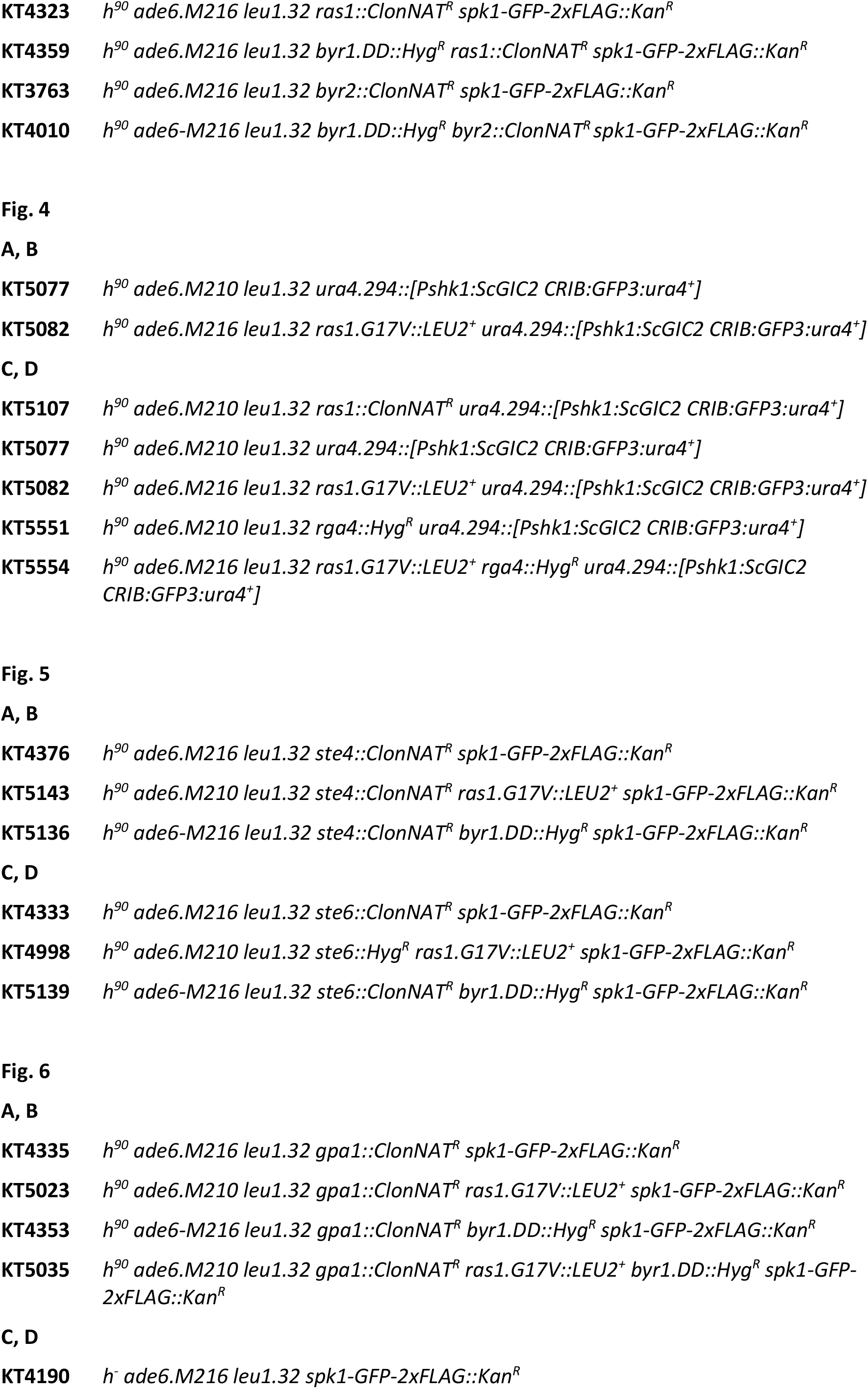
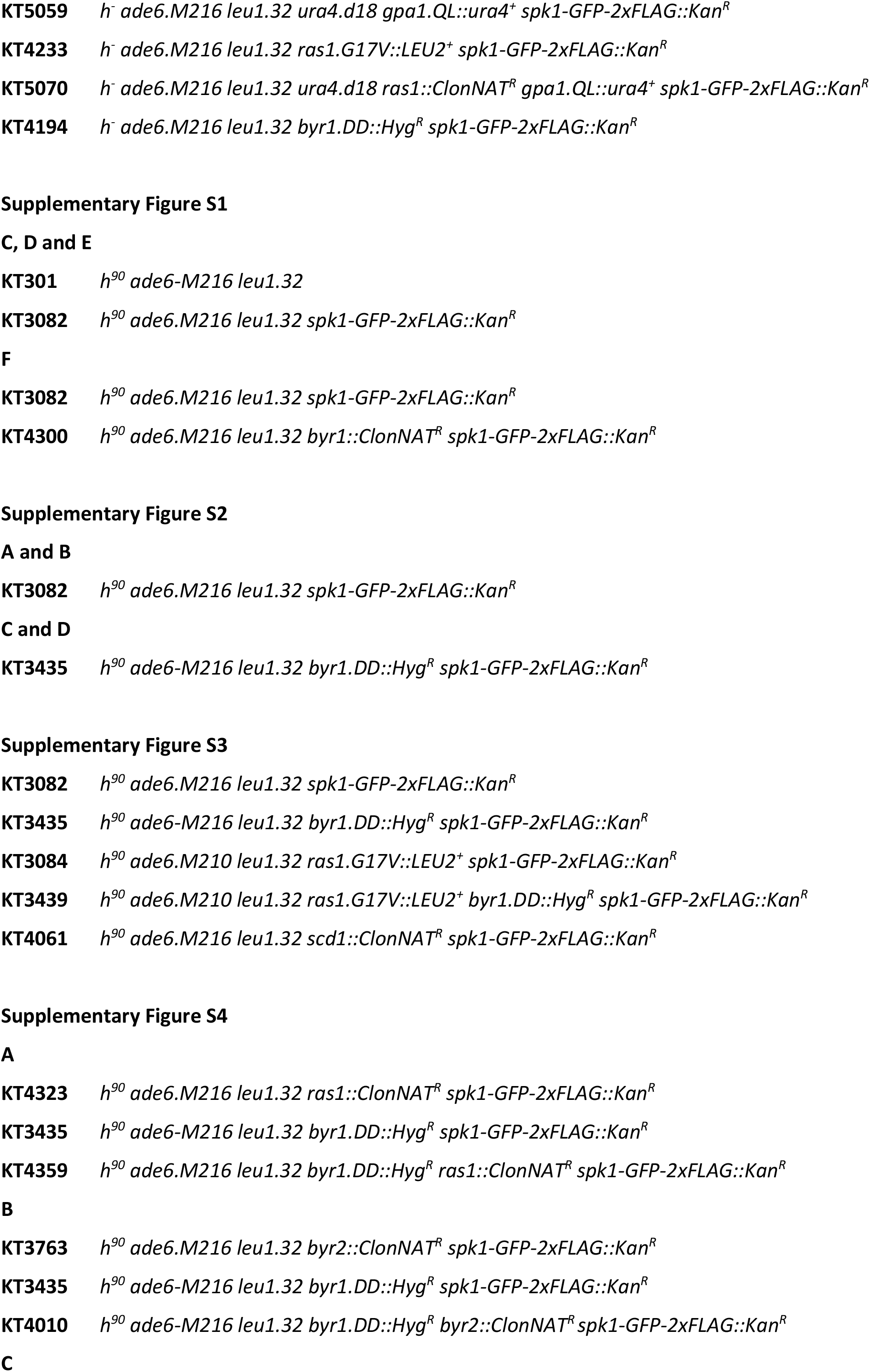
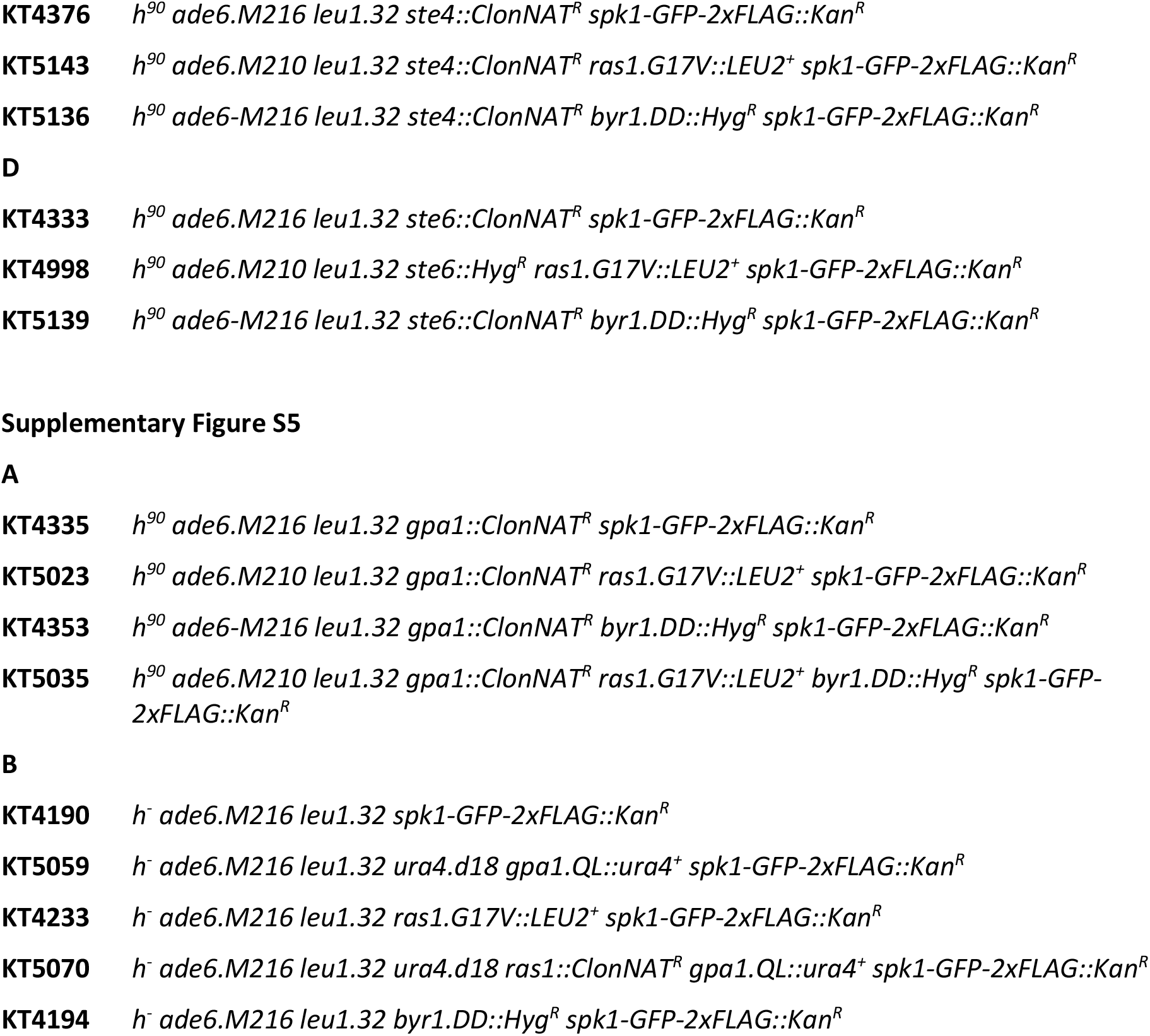
Strains used in this study

